# Identification of differentially expressed genes and enriched pathways in inflammatory bowel disease using bioinformatics and next generation sequencing data analysis

**DOI:** 10.1101/2023.06.01.543204

**Authors:** Basavaraj Vastrad, Chanabasayya Vastrad

## Abstract

Inflammatory bowel disease (IBD) is the most common chronic digestive disorders and inflammation in the gastrointestinal tract globally that is characterized by episodes of abdominal pain, diarrhea, bloody stools and weight loss. However, the pathophysiologic mechanisms of IBD have not been thoroughly investigated. To explore potential targets for treatment of IBD, we reorganized and analyzed next generation sequencing (NGS) dataset (GSE186507). The R package DESeq2 tool was used to screen for differentially expressed genes (DEGs) between IBD and normal control samples. We used the g:Profiler database to perform Gene Ontology (GO) enrichment analysis and the REACTOME for pathway enrichment analysis. Protein-protein interaction (PPI) network construction and module analysis were performed to elucidate molecular mechanisms of DEGs and screen hub genes. Then miRNA-hub gene regulatory network and TF-hub gene regulatory network of these hub genes were visualized by Cytoscape. We also validated the identified hub genes via receiver operating characteristic (ROC) curve analysis. A total of 957 DEGs (478 up regulated genes and 479 down regulated genes) were detected in NGS dataset. And they were mainly enriched in the terms of multicellular organismal process, response to stimulus, GPCR ligand binding and immune system. Based on the data of PPI network, miRNA-hub gene regulatory network and TF-hub gene regulatory network the top hub genes were ranked, including IL7R, ERBB2, SMAD1, RPS26, TLE1, HNF4A, CDKN1A, SRPK1, H3C12 and SFN. In conclusion, the identified DEGs, particularly the hub genes, strengthen the understanding of the development and progression of IDB, and certain novel genes might be used as candidate target molecules to diagnose, monitor and treat IDB.

## Introduction

Inflammatory bowel disease (IBD) is a chronic digestive disorders and inflammation in the gastrointestinal tract characterized by episodes of abdominal pain, diarrhea, bloody stools, weight loss, and the influx of neutrophils and macrophages that produce cytokines, proteolytic enzymes, and free radicals that result in inflammation and ulceration [1]. There are two main types of IBD: Crohn’s disease and ulcerative colitis [2]. While it is mostly associated with intestinal strictures [3], fistulas [4], abscesses [5], malnutrition [6], autoimmune diseases [7], coagulation and fibrinolysis [8], gastrointestinal cancers [9], osteoporosis [10] and anemia [11]. The prevalence of IBD appears to be increasing worldwide, with the highest rates of increase seen in developing countries [12]. According to latest estimates, the worldwide prevalence of IBD is approximately 0.5-0.7% of the population, or 5-7 cases per 1,000 people [13]. This might be due to a combination of genetic [14], environmental [15], and lifestyle factors [16]. However, few effective therapies of IBD are still a clinical challenge [17].

The molecular mechanisms of IBD and progression remain unclear. It is therefore critical to identify novel genes and signaling pathways that are associated with IBD and patient prognosis, which might not only help to elucidate the underlying molecular mechanisms involved, but also to discover new diagnostic markers and therapeutic targets. For example, studies have found that genes include IL2RA and IL2RB [18], HIF-1α [19], IL-10 [20], NOD2 [21] and IL-23 [22], and signaling pathways include JAK-STAT signaling pathway [23], AhR/Nrf2/NQO1 signaling pathway [24], TAT-3 and Rac-1 signaling pathway [25], Wnt and Notch signaling pathways [26], and interleukin-23/Th17 signaling pathways [27] were responsible for IBD progression. However, because of heterogeneity, there is still an urgent need to identify new biomarkers that might aid the diagnosis and treatment of IBD.

Next generation sequencing (NGS), which allow the investigation of gene expression in a high throughput manner with high sensitivity, specificity and repeatability [28]. A significant amount of data has been produced via the use of NGS data has been uploaded and stored in public databases. Previous investigation concerning IBD gene expression profiling has identified hundreds of DEGs [29]. One potential interpretation is that unidentified genes might partially contribute to the missing heritability. Therefore, there are still various related biomarkers to be identified, which will help us better understand the molecular pathogenesis of IBD and facilitate the discovery of novel diagnostic and prognostic biomarkers or therapeutic target.

Bioinformatics analysis is an important strategy for the comprehensive analysis of huge databases, including arduous genetic information. In our investigation, we used sophisticated bioinformatics methods to screen potential biomarkers that might be useful for IBD. The Gene Expression Omnibus (GEO) [https://www.ncbi.nlm.nih.gov/geo/] database [30] is an open database that allows researchers to select appropriate NGS data. In our investigation, we obtained three NGS dataset from the GEO (GSE186507) [31] and searched for DEGs using DESeq2 package in R software [32]. We then performed gene ontology (GO) and REACTOME pathway enrichment analyses of the identified DEGs using the g:Profiler database. Protein-protein interaction (PPI) network was constructed using IMEX interactome database and visualized using Cytoscape. Conduct module analyses of the PPI network were performed using PEWCC. MiRNA-hub gene regulatory network and TF-hub gene regulatory network construction were carried out according to the screened hub genes. The screened key genes were validated using a receiver operating characteristic (ROC) curve analysis. Collectively, the findings of the current investigation highlighted essential genes and pathways that might contribute to the pathology of IBD. These might provide a basis for the advancement of future diagnostic, prognostic and therapeutic tools for IBD.

## Materials and Methods

### Next generation sequencing data source

(GSE186507) [31] NGS dataset was downloaded from the GEO database and based on a GPL16791 Illumina HiSeq 2500 (Homo sapiens). The dataset contained 1016 blood samples, including 807 IBD samples and 209 normal control samples.

### Identification of DEGs

The R package DESeq2 was then used to identify the DEGs among the IBD and normal control groups in NGS dataset. The Benjamini and Hochberg (BH) method was carried out to adjust P value to reduce the false positive error [33]. The cutoff criteria for determining DEGs were |log2 fold change (FC)| > 0.196 for up regulated genes, |log2 fold change (FC)| < -0.2941 for down regulated genes and adjusted P value < 0.05, which were visualized as Volcano plots and heat map plots. ggplot2 and gplot packages of R software was applied to generate Volcano plots and heat map plots

### GO and pathway enrichment analyses of DEGs

Functional enrichment analysis of DEGs was performed by the g:Profiler (http://biit.cs.ut.ee/gprofiler/) [34]. The g:Profiler was used to perform the Gene Ontology (GO, http://www.geneontology.org) [35] enrichment analysis and REACTOME (https://reactome.org/) [36] pathway enrichment analysis. Through GO enrichment analysis, the biological processes (BP), cellular components (CC) and molecular functions (MF) involved in various genes were interpreted. REACTOME pathway enrichment analysis was conducted to examine the role of DEGs in different signal transduction pathways in the human body. A adjusted P value < 0.05 in both GO and REACTOME pathway enrichment analyses was set as the threshold for significant enrichment.

### Construction of the PPI network and module analysis

DEGs were uploaded to The International Molecular Exchange Consortium (IMEX interactome, http://www.imexconsortium.org/) [37] database to analyze interactions among the proteins encoded by the identified DEGs. Results with a minimum interaction score of 0.4 were visualized using Cytoscape (http://www.cytoscape.org/) [38]. The PPI network for hub genes was computed with the maximal degree [39], betweenness [40], stress [41] and closeness [42] methods and Network Analyzer. PEWCC [43] plugin, the PPI network was divided into two modules.

### Construction of the miRNA-hub gene regulatory network

miRNet database (https://www.mirnet.ca/) [44] is a bioinformatics platform for predicting miRNAs and miRNA hub gene pairs. In the present investigation, the targets of the miRNAs were predicted using 14 databases: TarBase, miRTarBase, miRecords, miRanda (S mansoni only), miR2Disease, HMDD, PhenomiR, SM2miR, PharmacomiR, EpimiR, starBase, TransmiR, ADmiRE, and TAM 2.0. The screening criterion was that the miRNA target exists in the 14 databases concurrently. The miRNA hub gene regulatory network was depicted and visualized using Cytoscape software [38].

### Construction of the TF-hub gene regulatory network

NetworkAnalyst database (https://www.networkanalyst.ca/) [45] is a bioinformatics platform for predicting TFs and TF hub gene pairs. In the present investigation, the targets of the TFs were predicted using CHEA TF database. The screening criterion was that the TF target exists in the CHEA database. The TF hub gene regulatory network was depicted and visualized using Cytoscape software [38].

### Receiver operating characteristic curve (ROC) analysis

ROC curve analyses to determine the specificity, sensitivity, likelihood ratios, positive predictive values, and negative predictive values for all possible thresholds of the ROC curve were performed using the pROC package in R statistical software [46]. Used one data as a training dataset and other data as a validation dataset iteratively. The receiver operator characteristic curves were plotted and area under curve (AUC) was calculated separately to evaluate the diagnostic value of the hub gene. AUCL>L0.9 indicated that the model had a good fitting effect.

## Results

### Identification of DEGs

NGS dataset GSE186507 was downloaded from the GEO database and analyzed using R packages (DESeq2, ggplot2 and gplot). The DEGs were screened by “DESeq2” package (|log2 fold change (FC)| > 0.196 for up regulated genes, |log2 fold change (FC)| < -0.2941 for down regulated genes and adjusted P value < 0.05). The GSE186507 dataset contained total 957 DEGs, including 478 up regulated genes and 479 down regulated genes (Table 1). Volcano plots were generated to visualize fold changes of the DEGs (Fig.1). The heatmap of the DEGs is shown in Fig. 2.

**Table 1.**
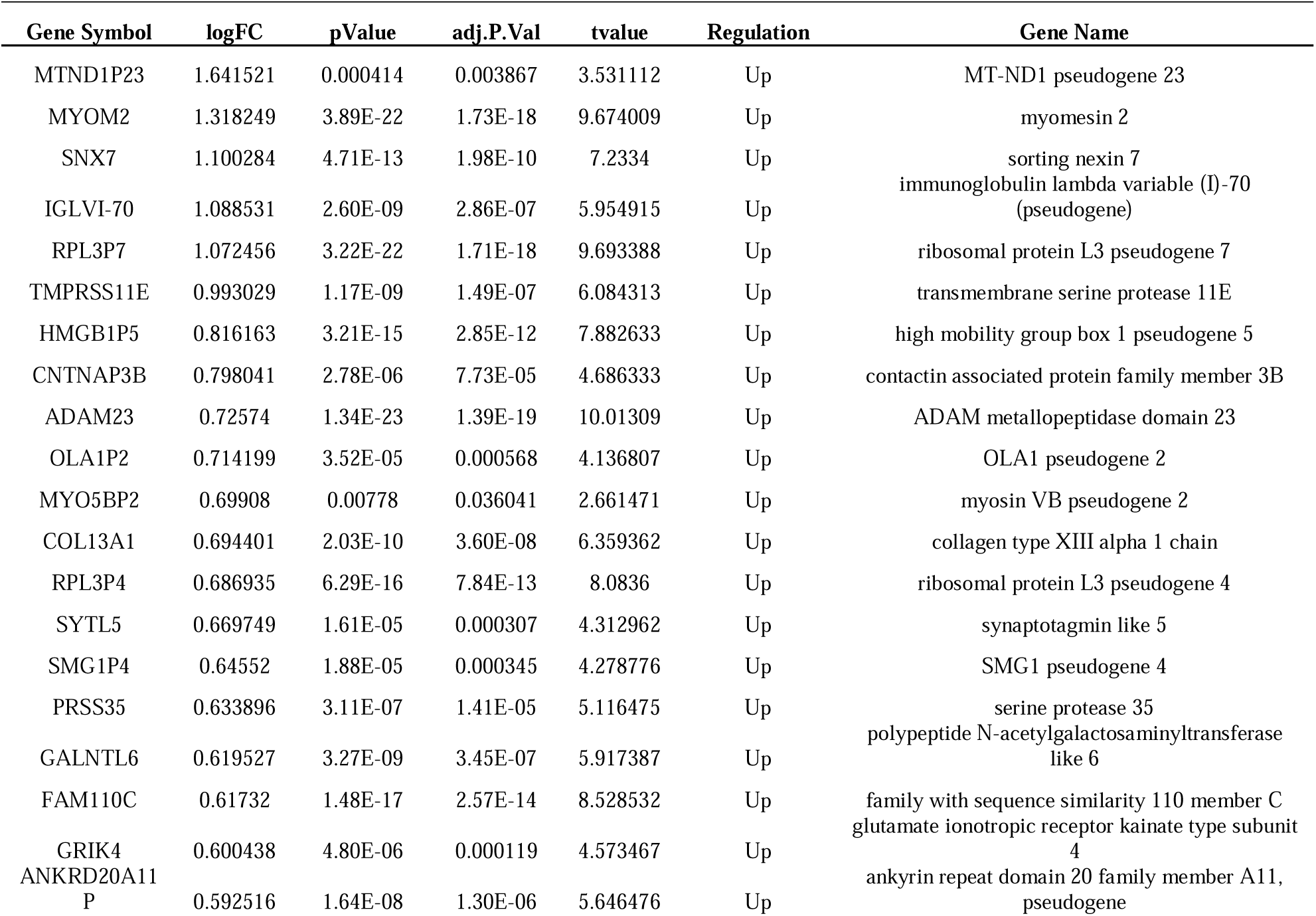
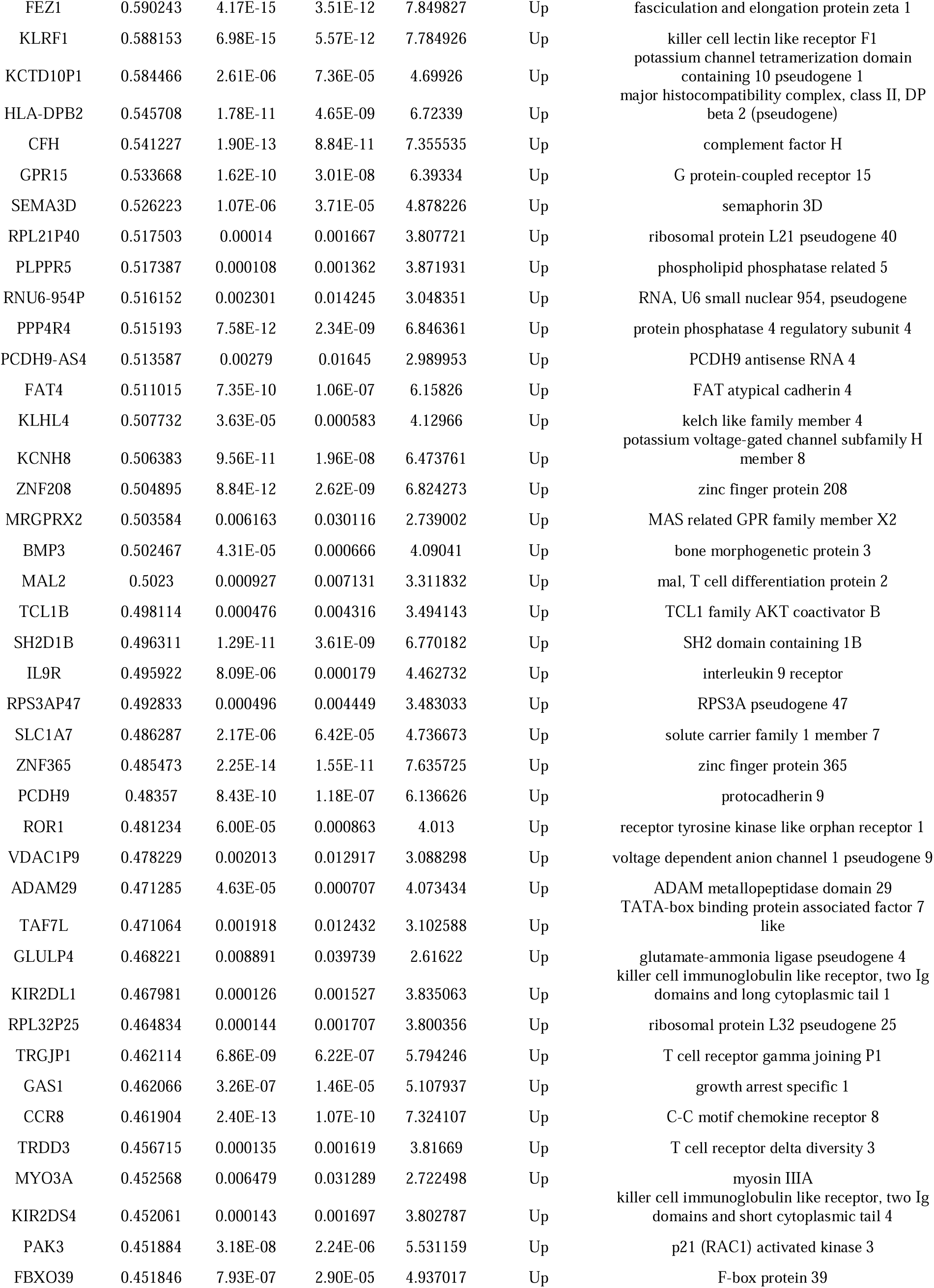
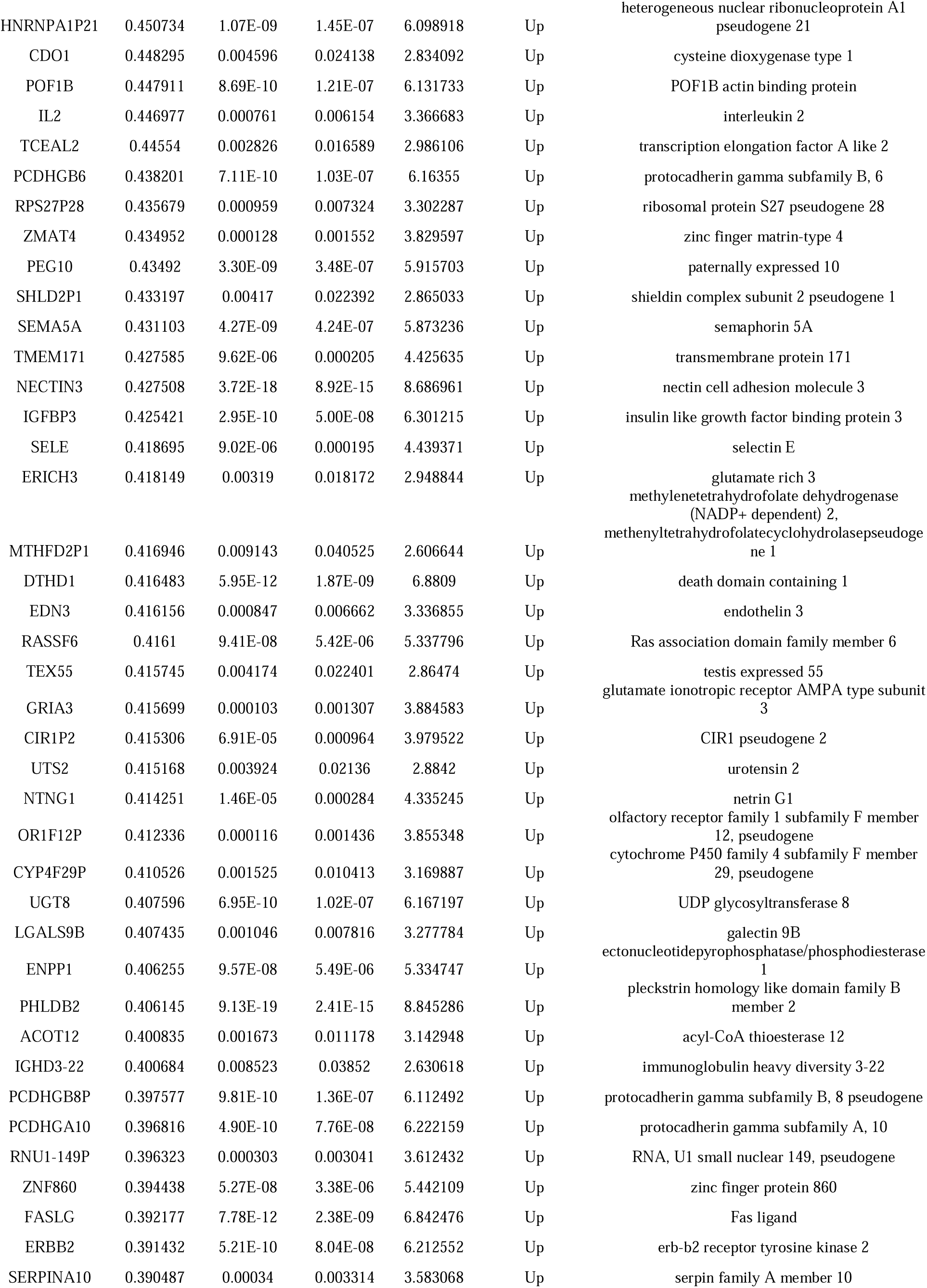
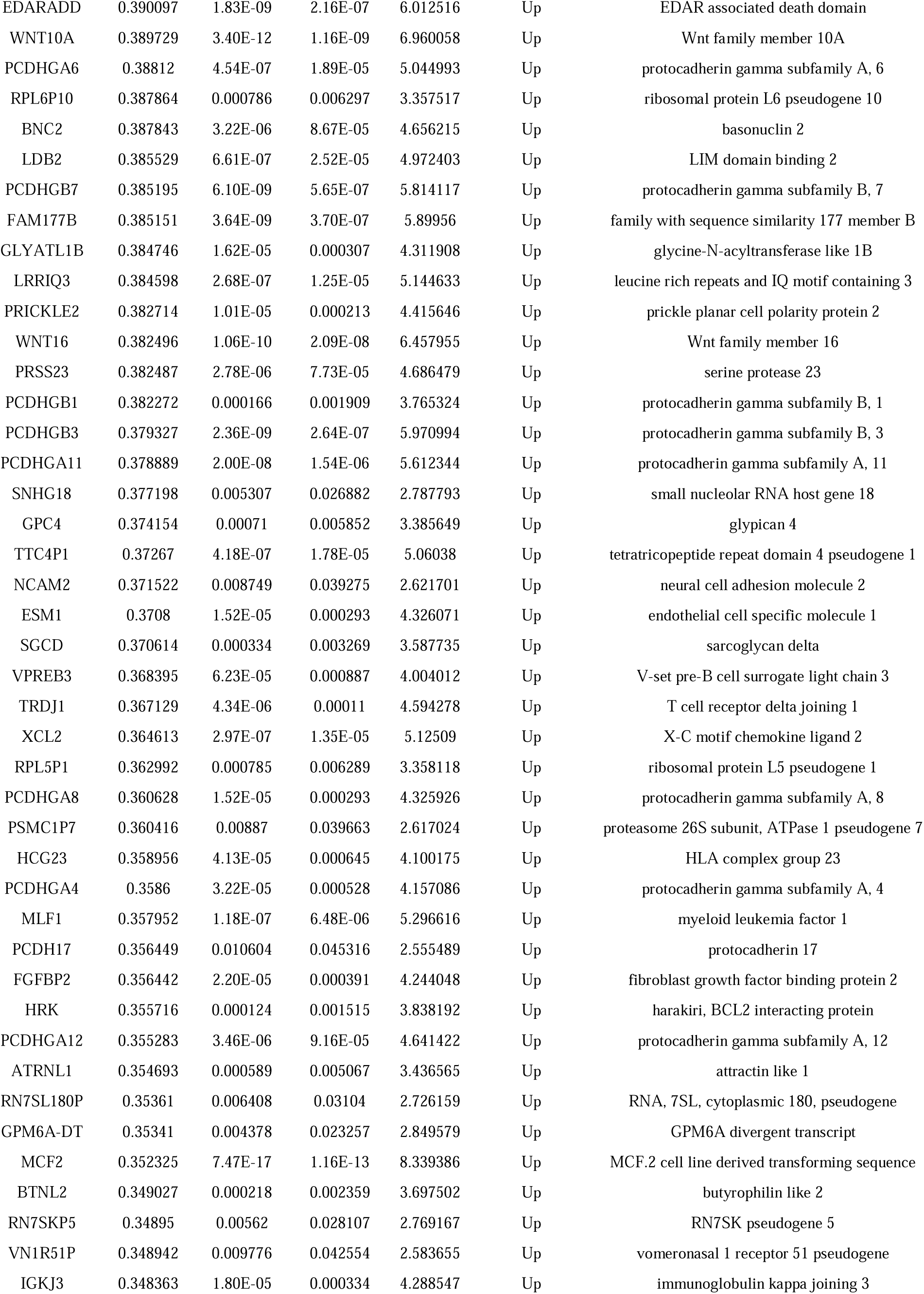
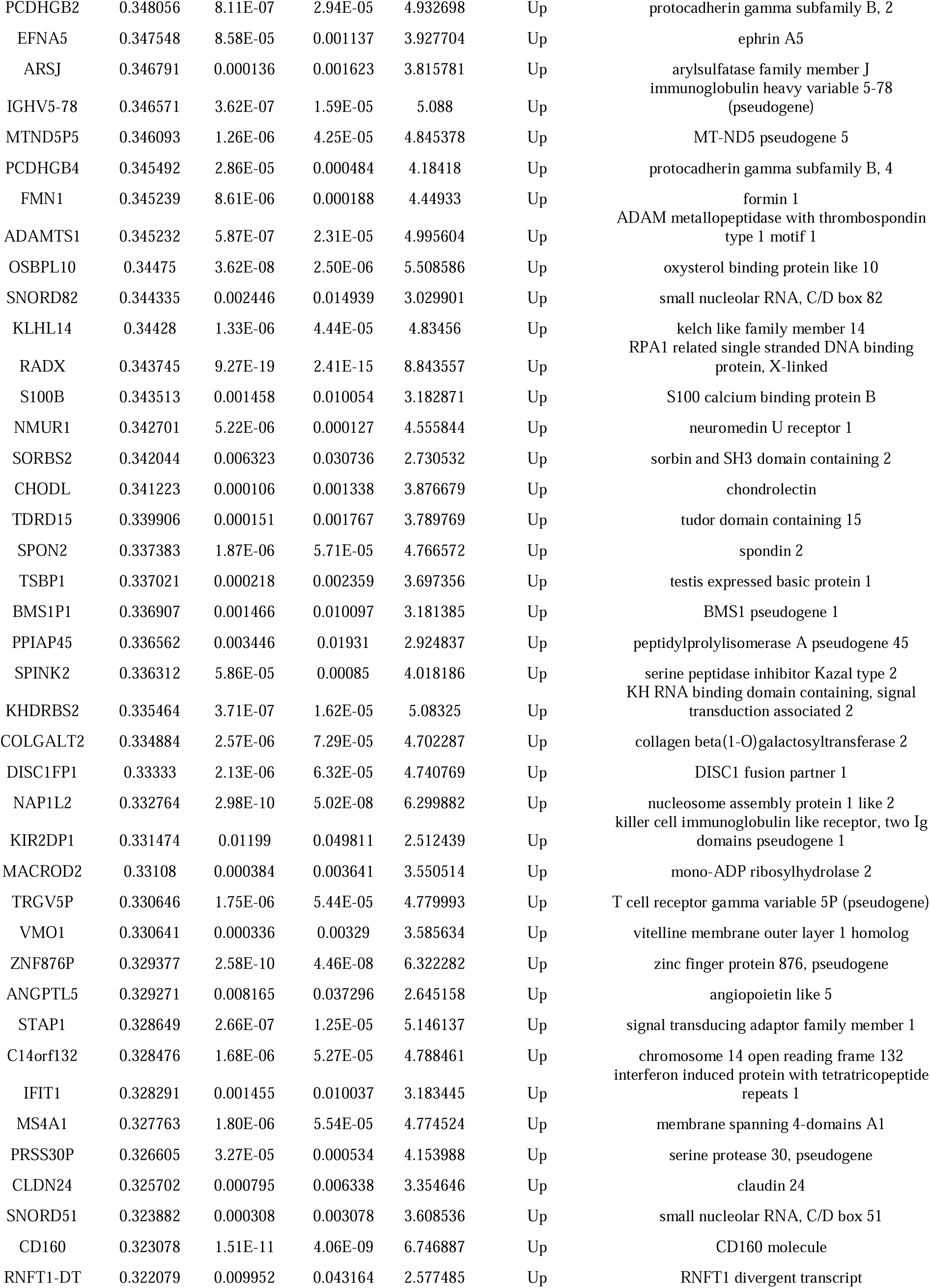
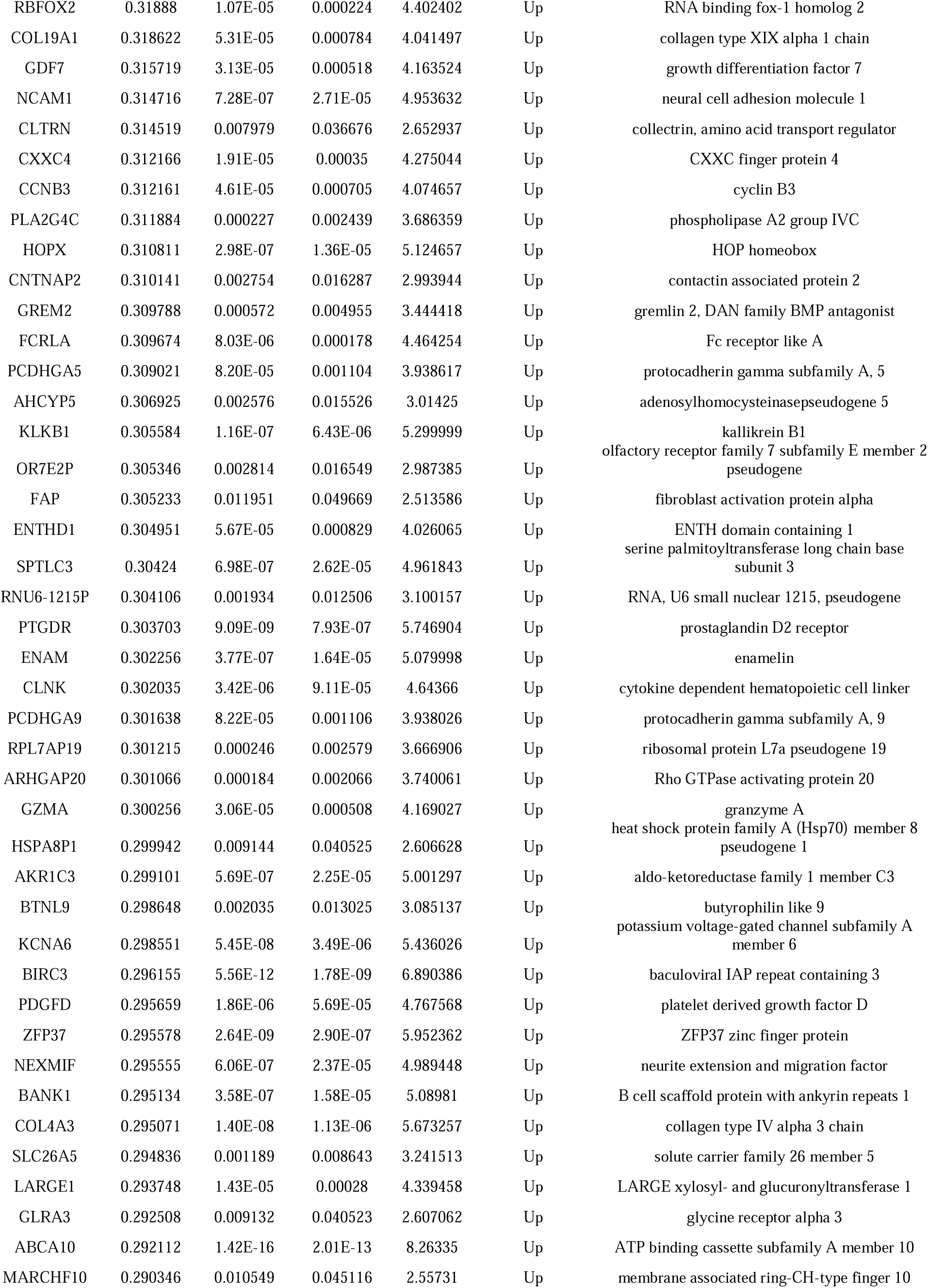
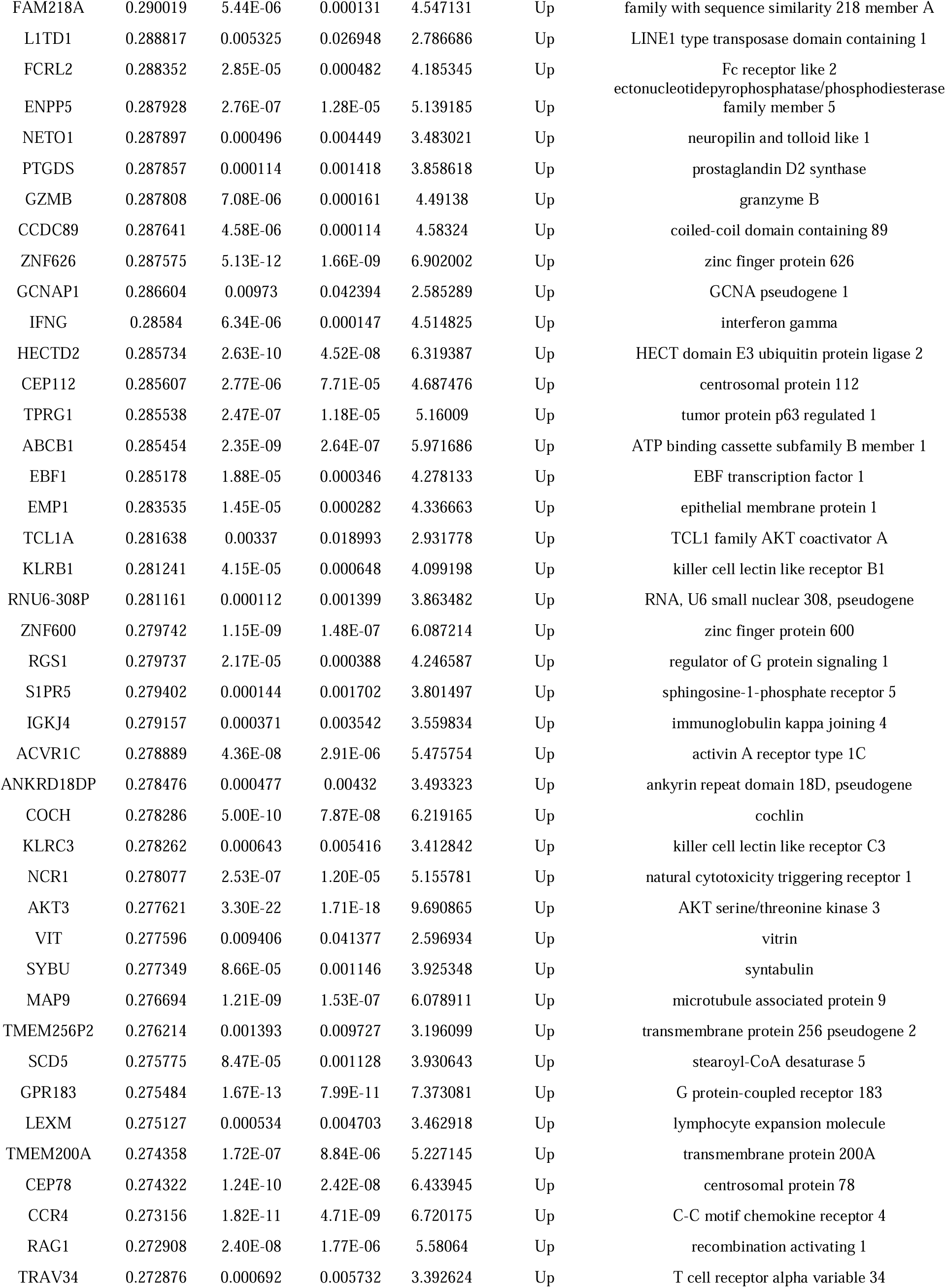
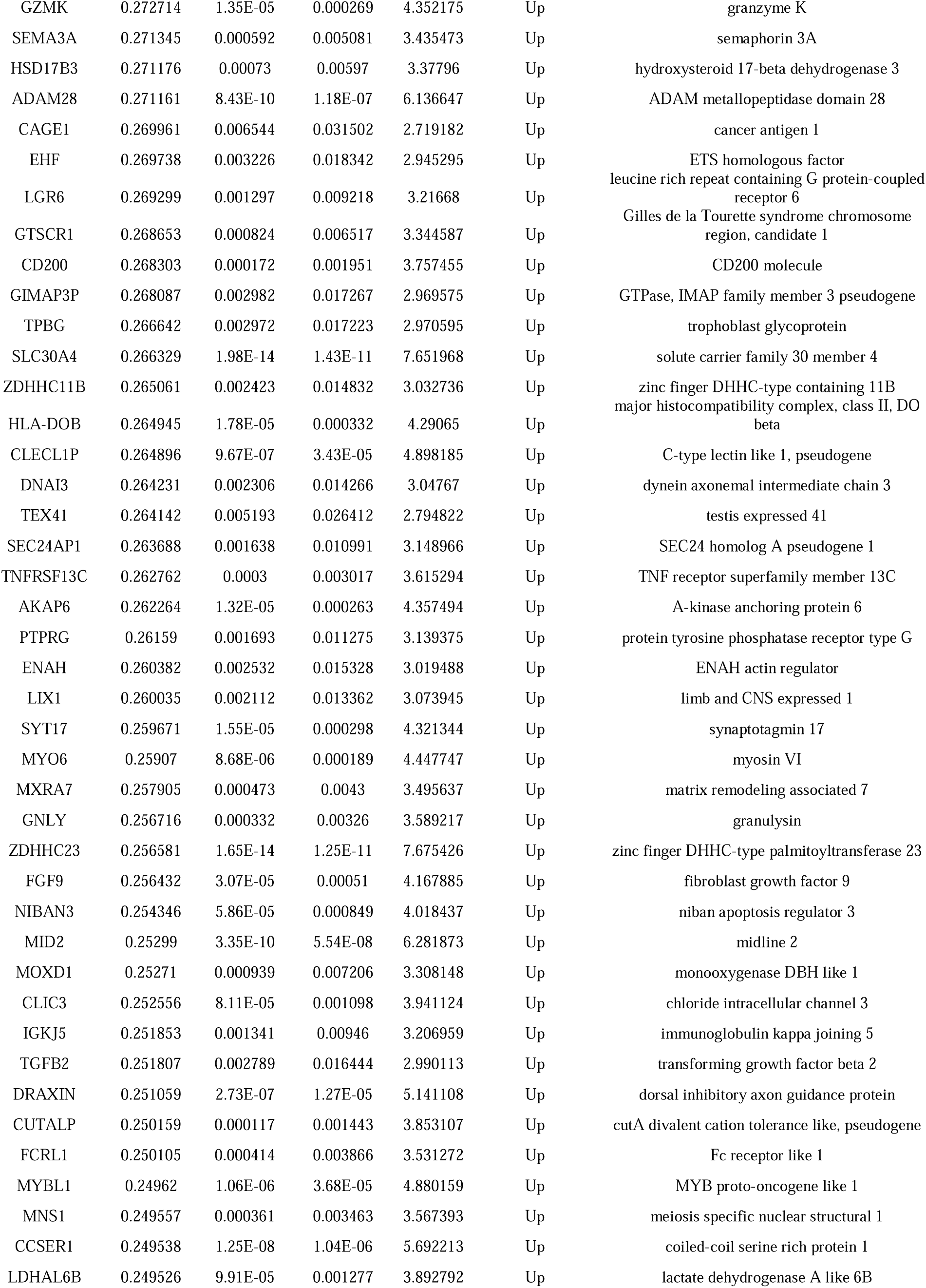
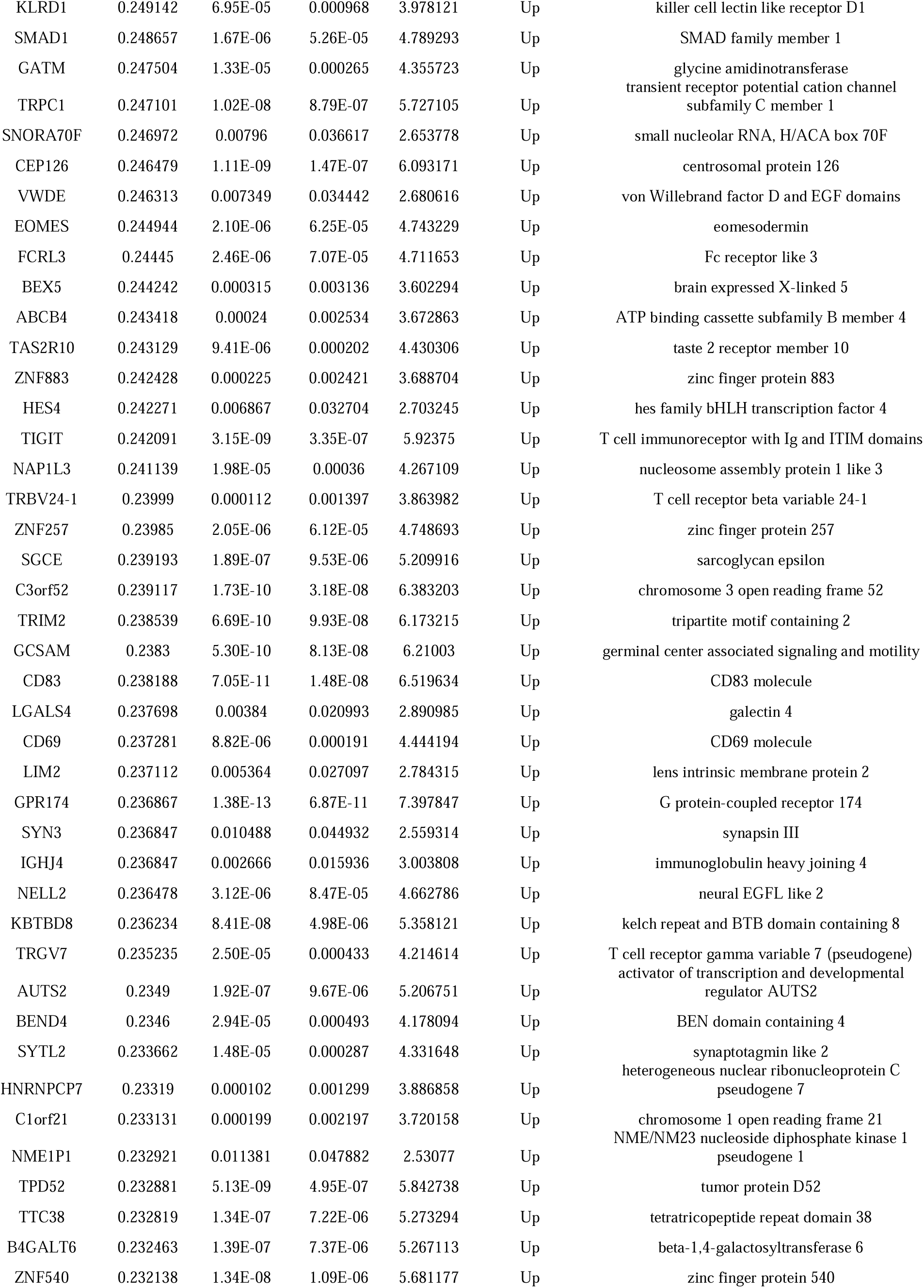
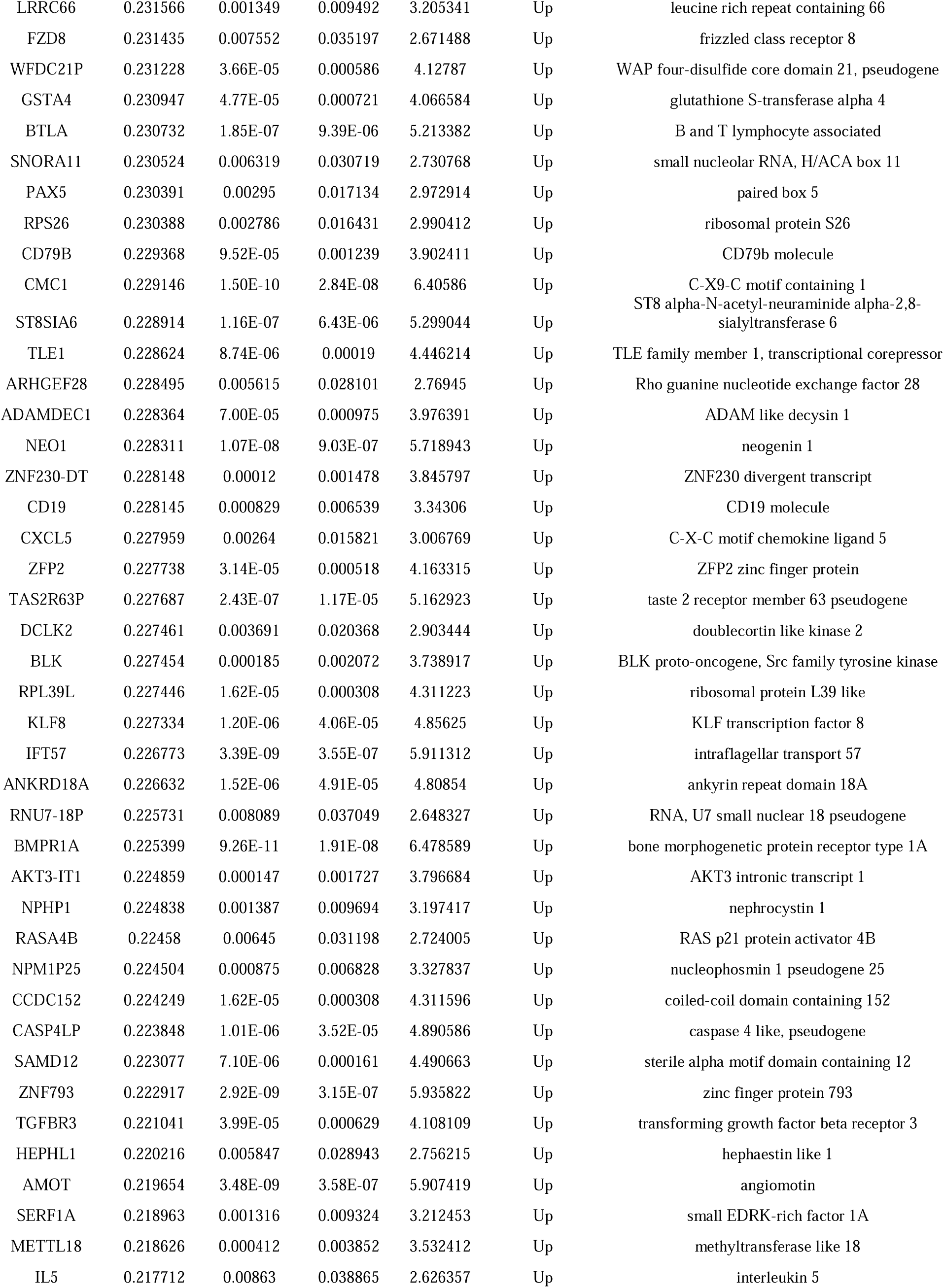
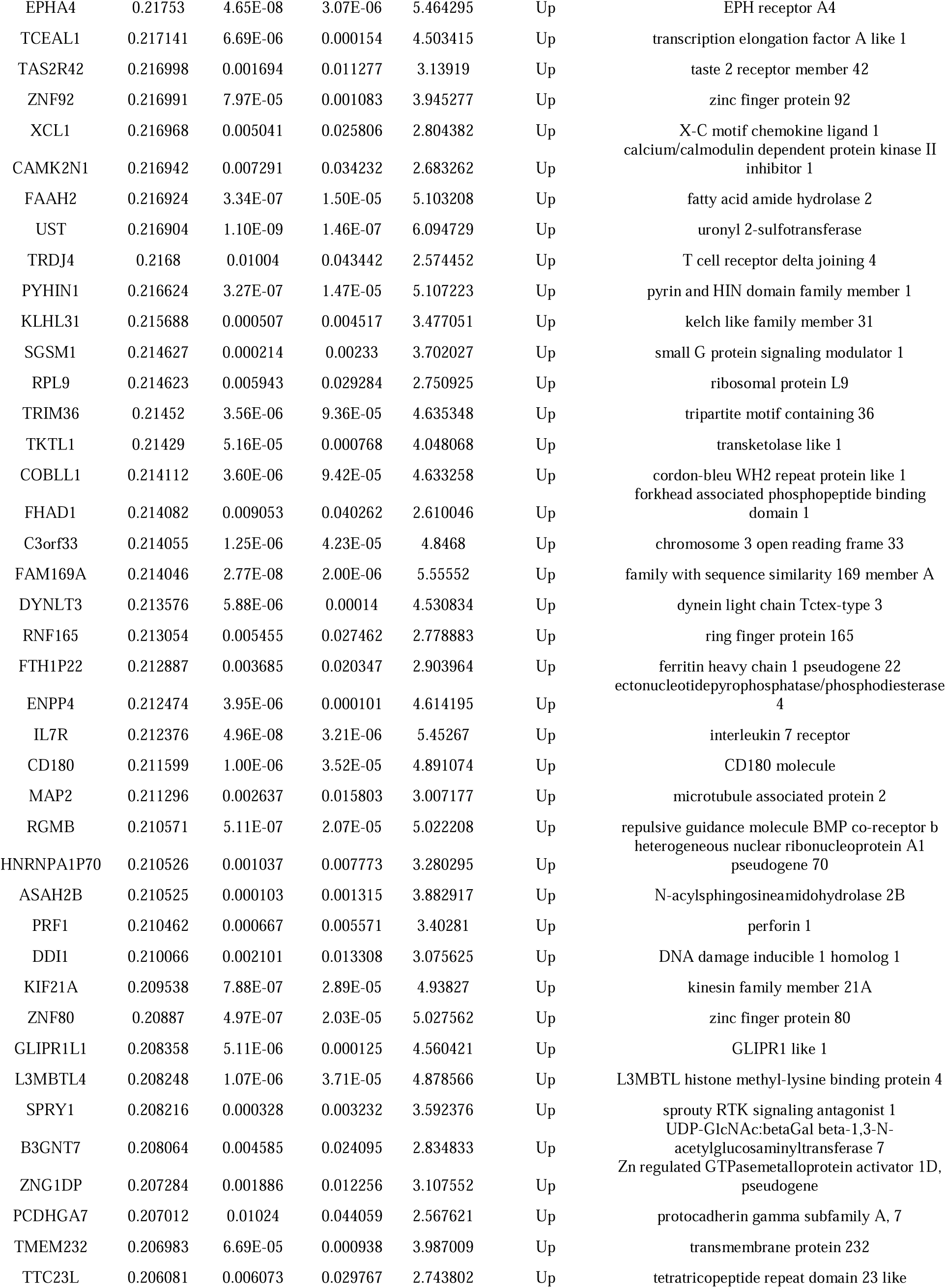
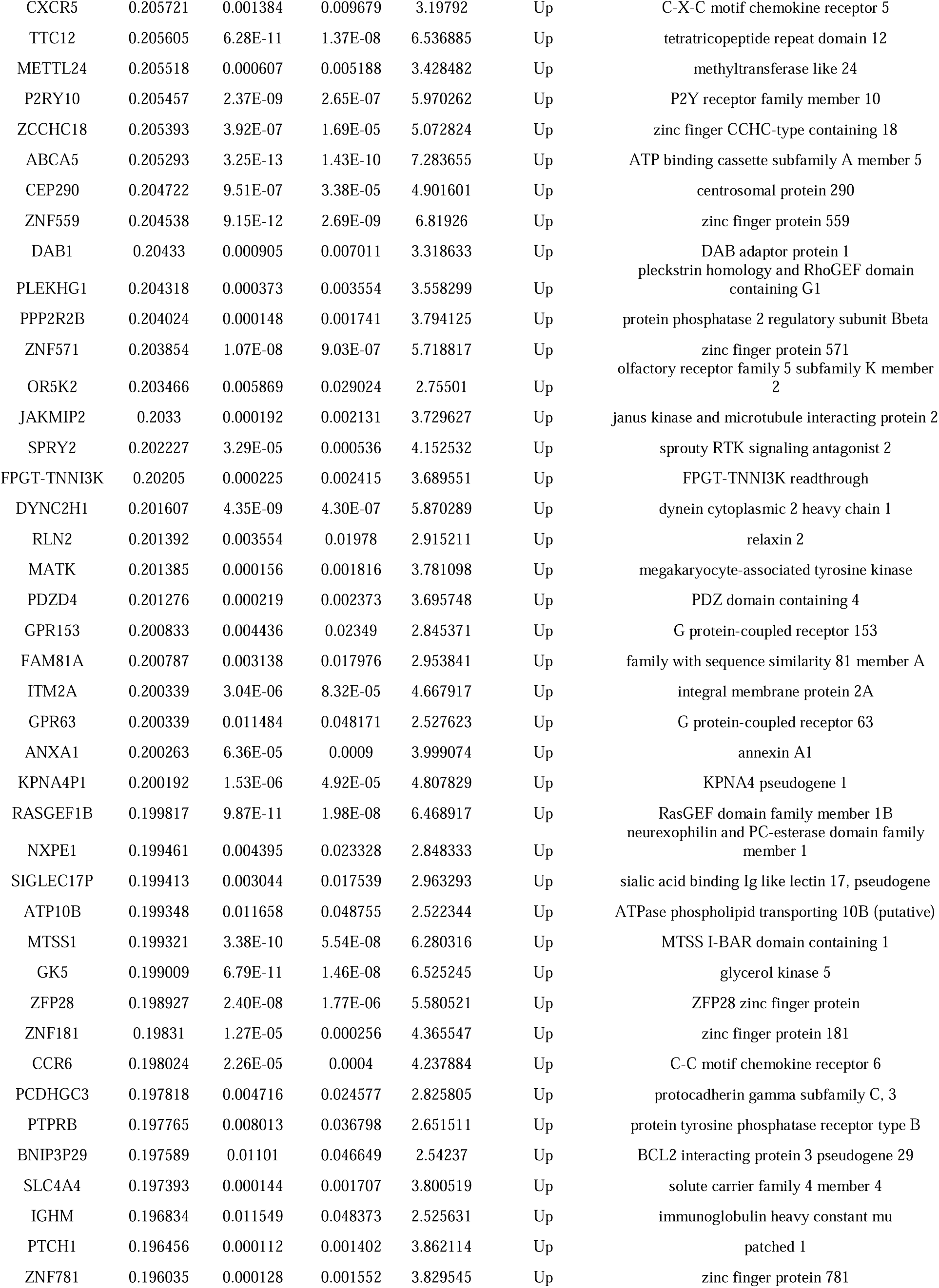
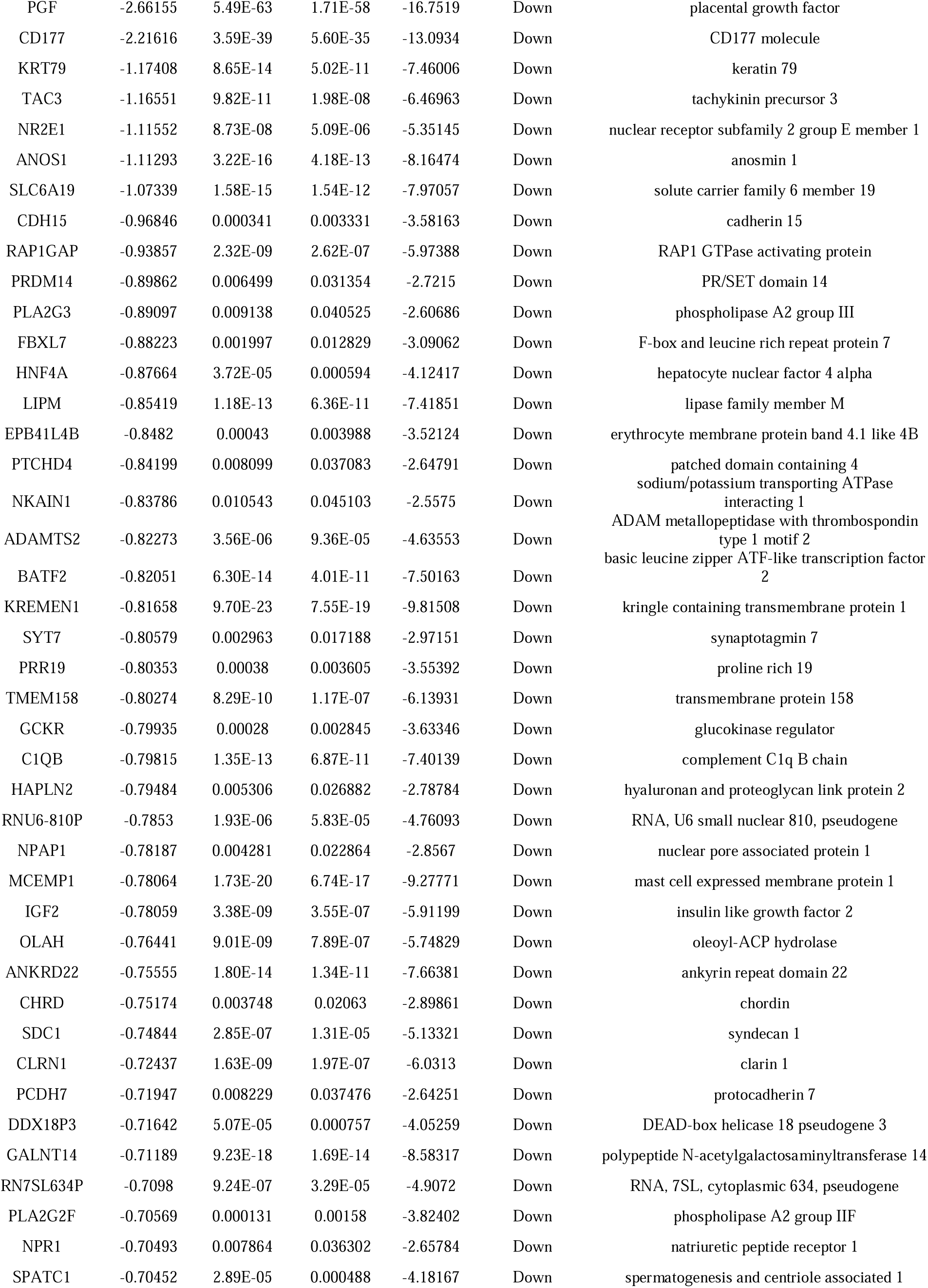
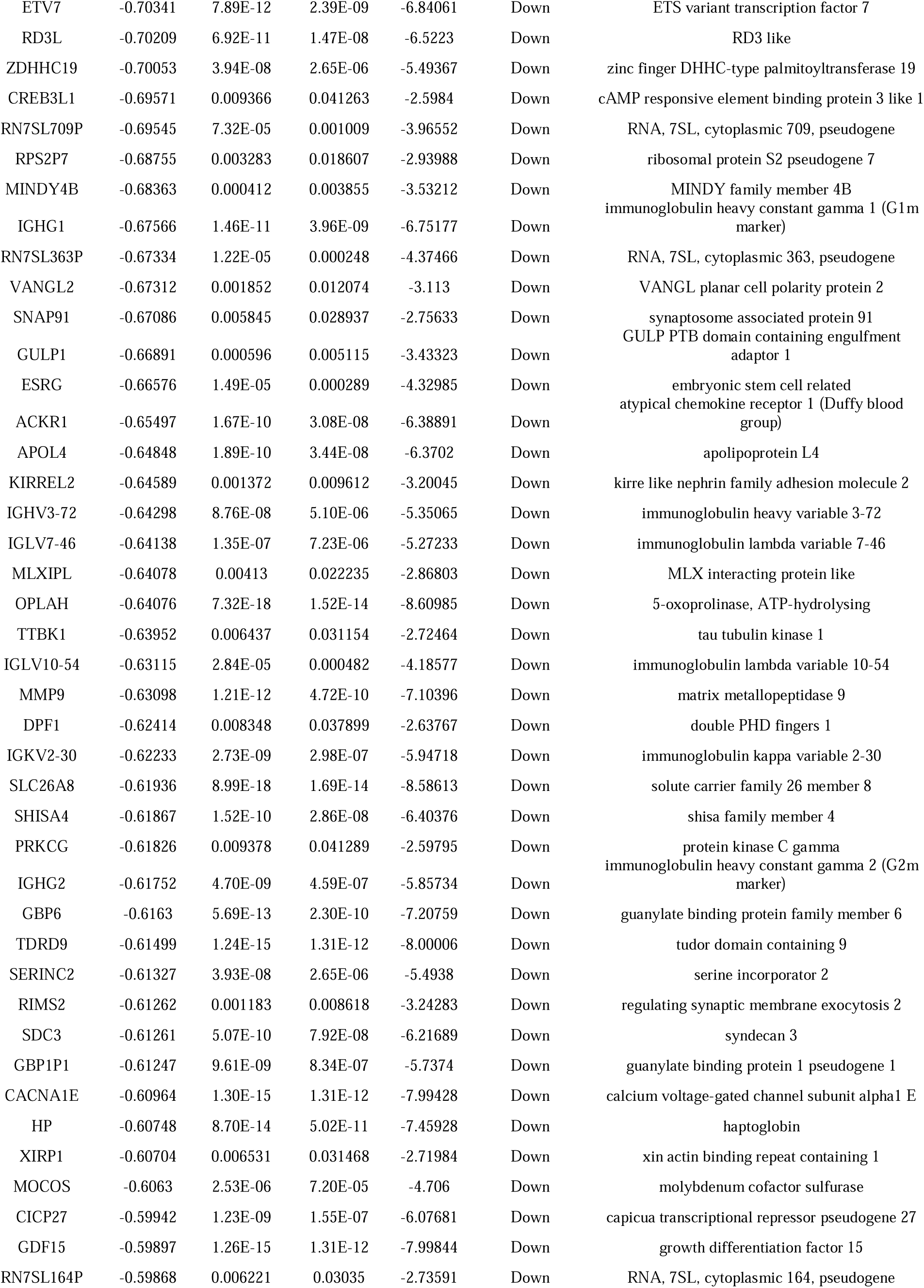
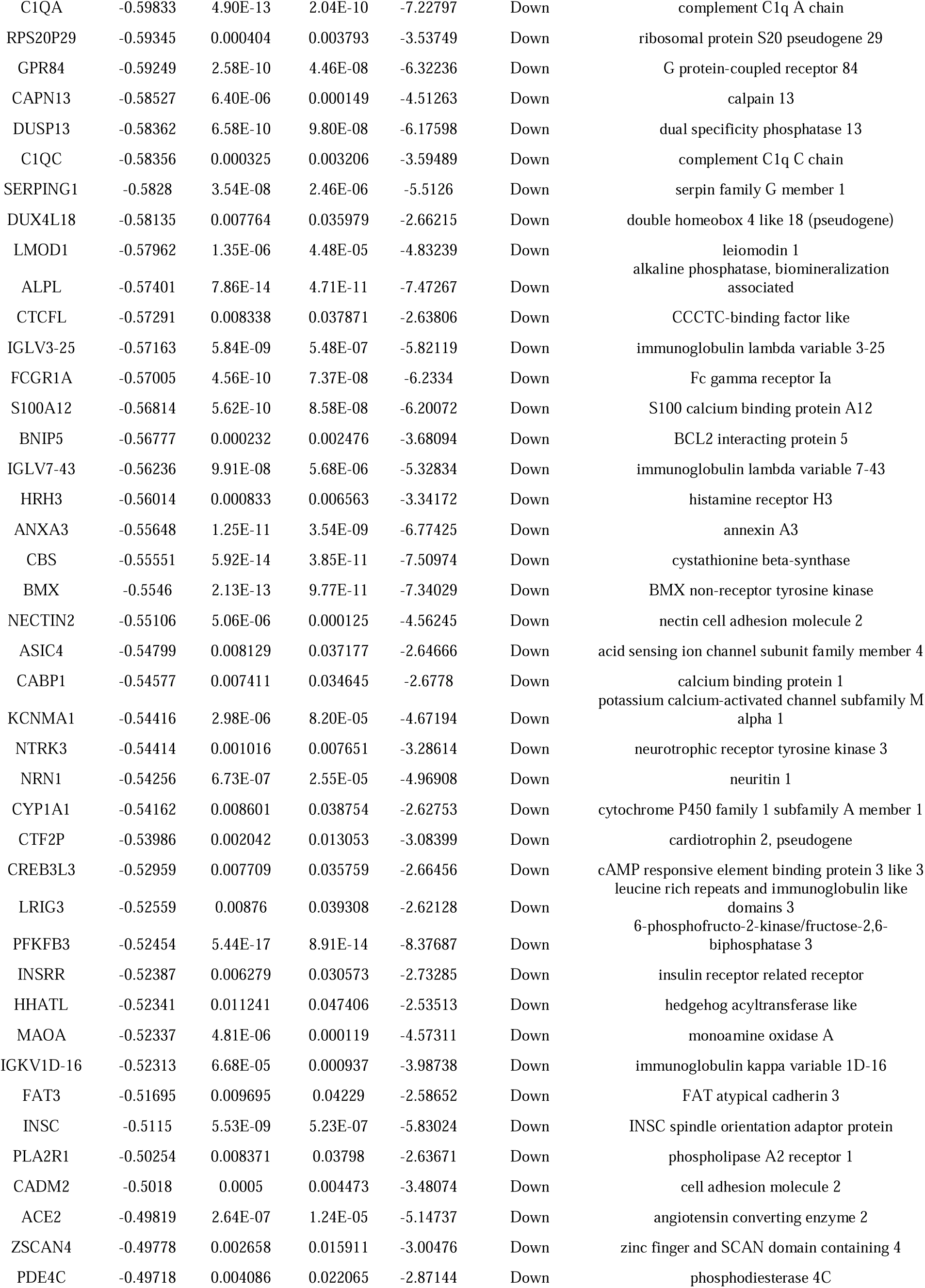
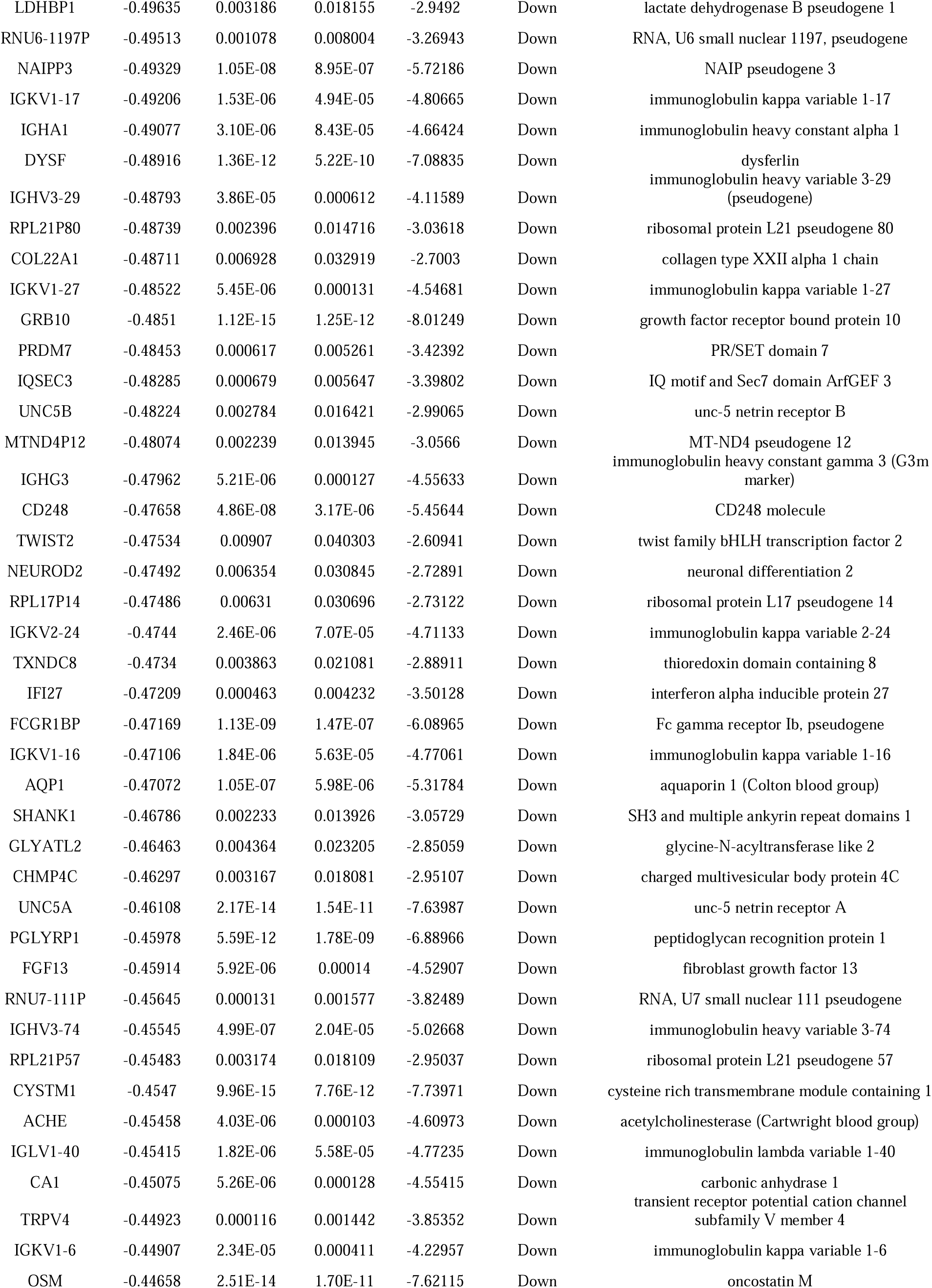
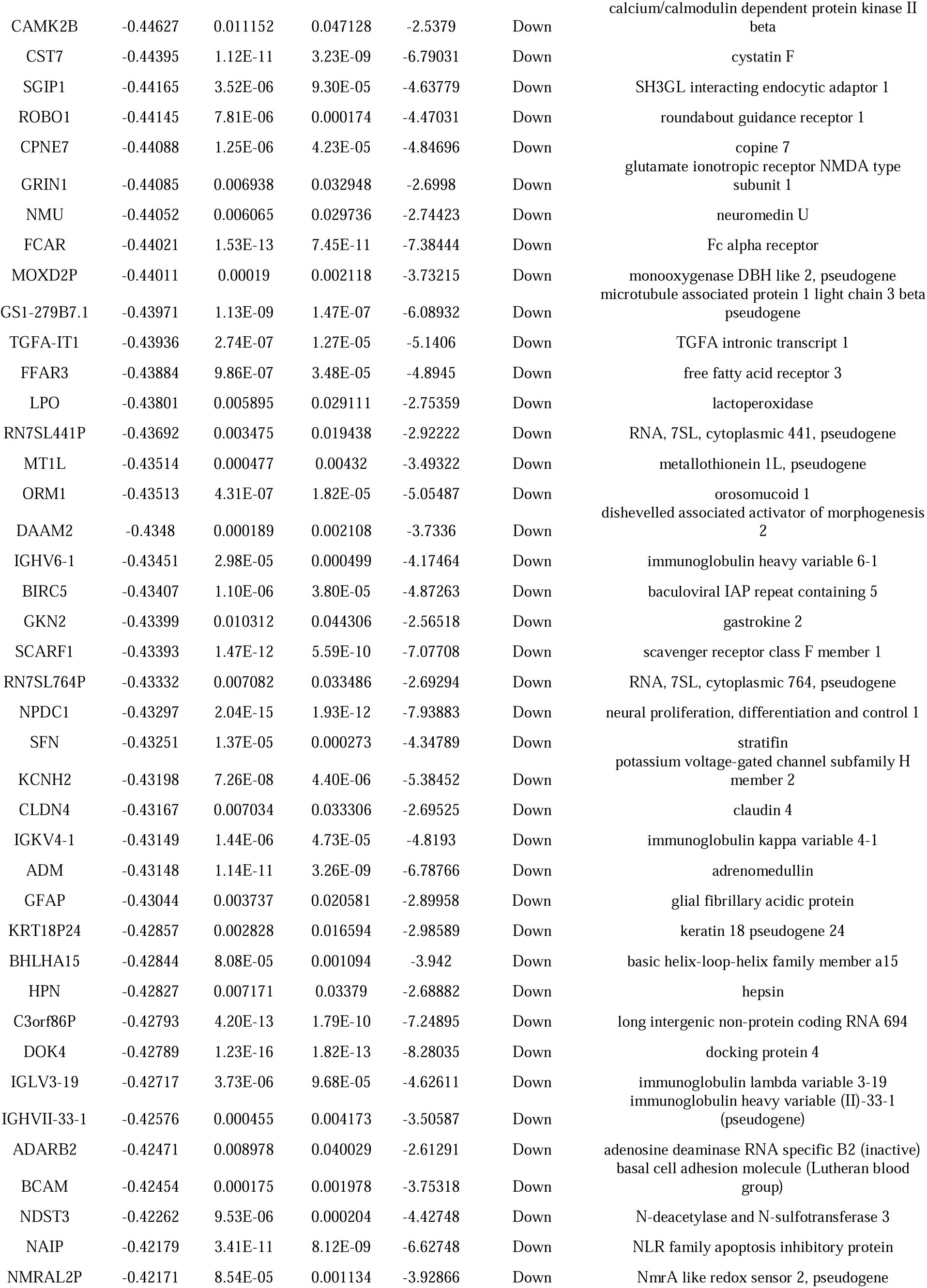
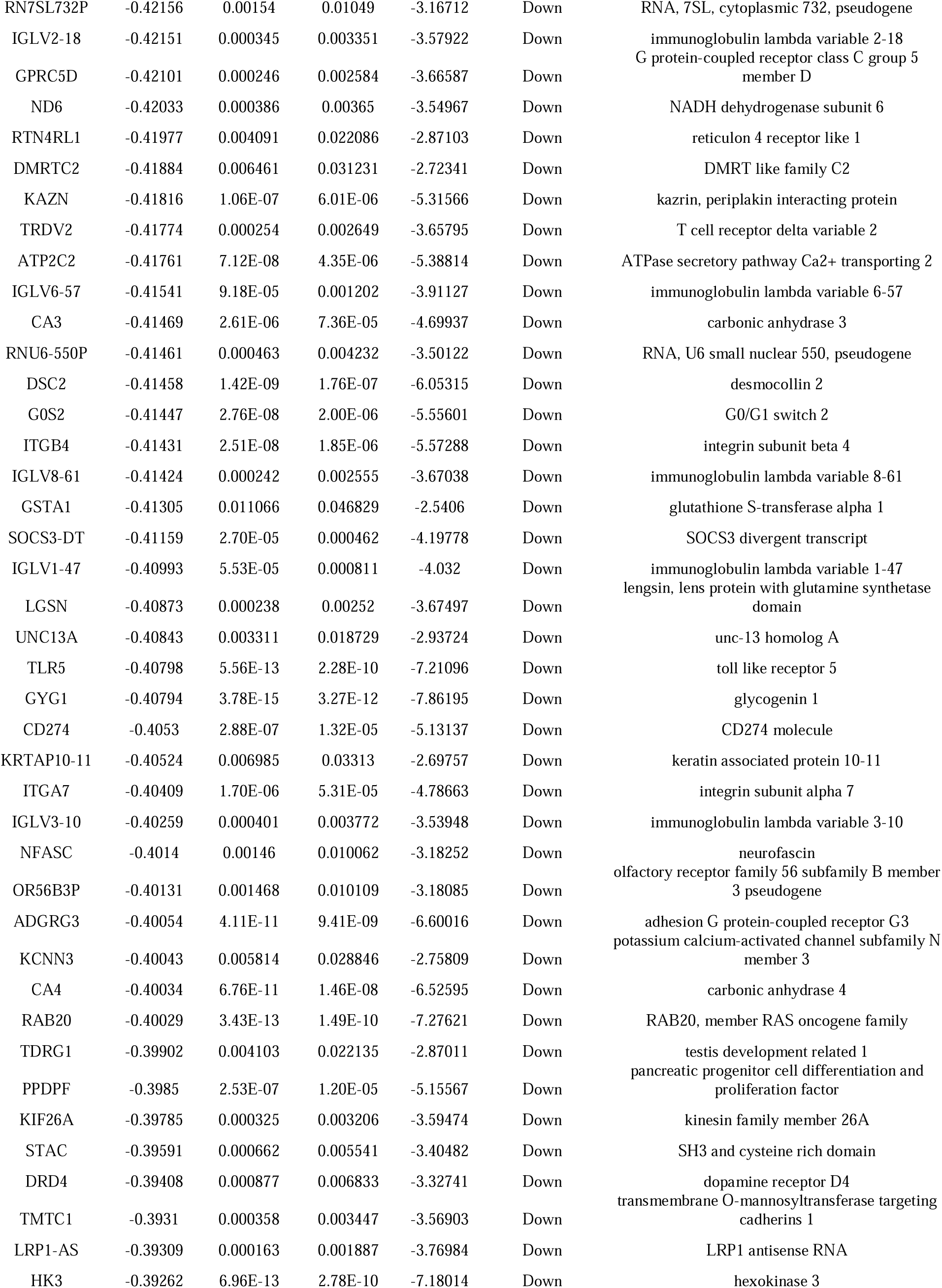
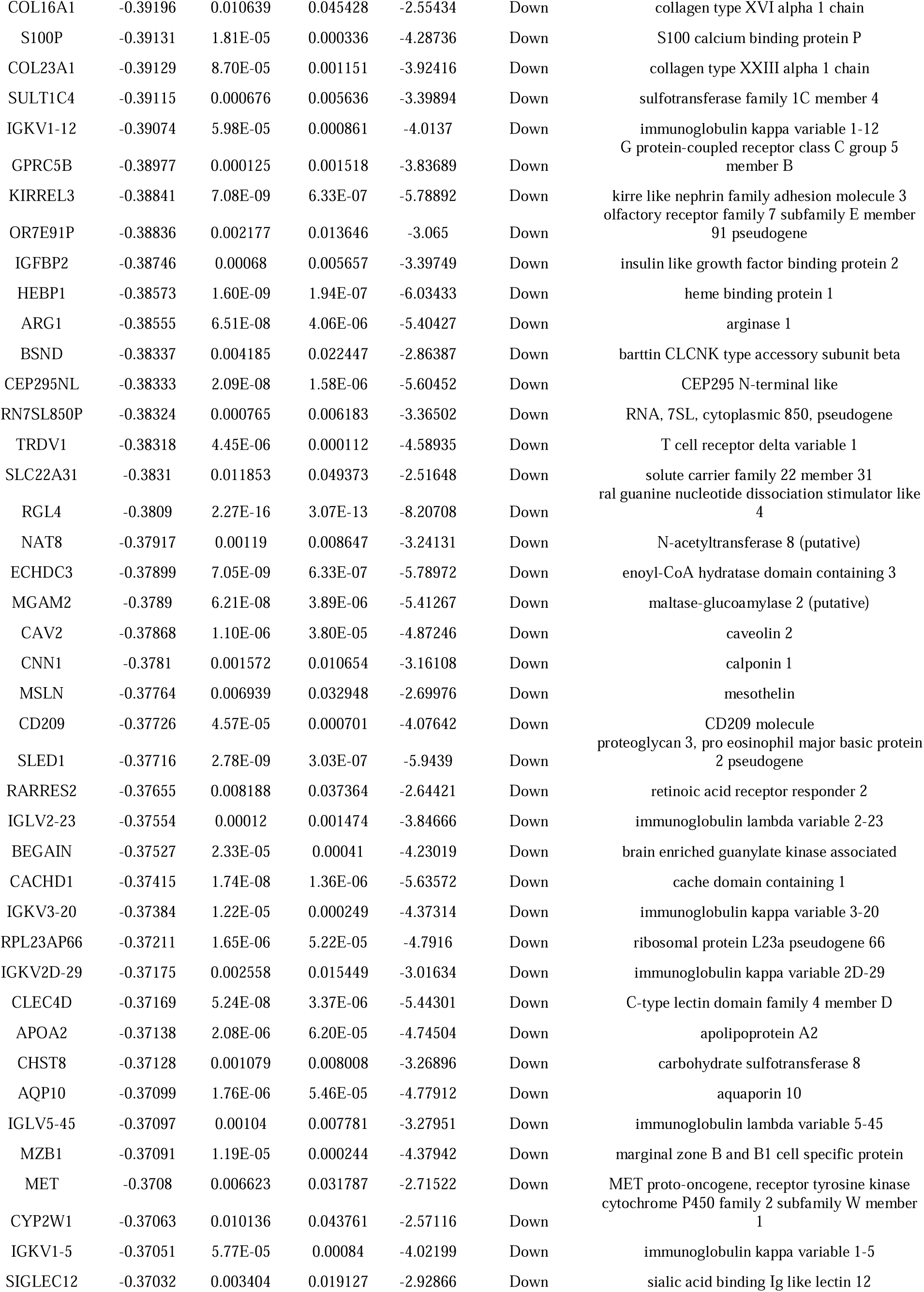
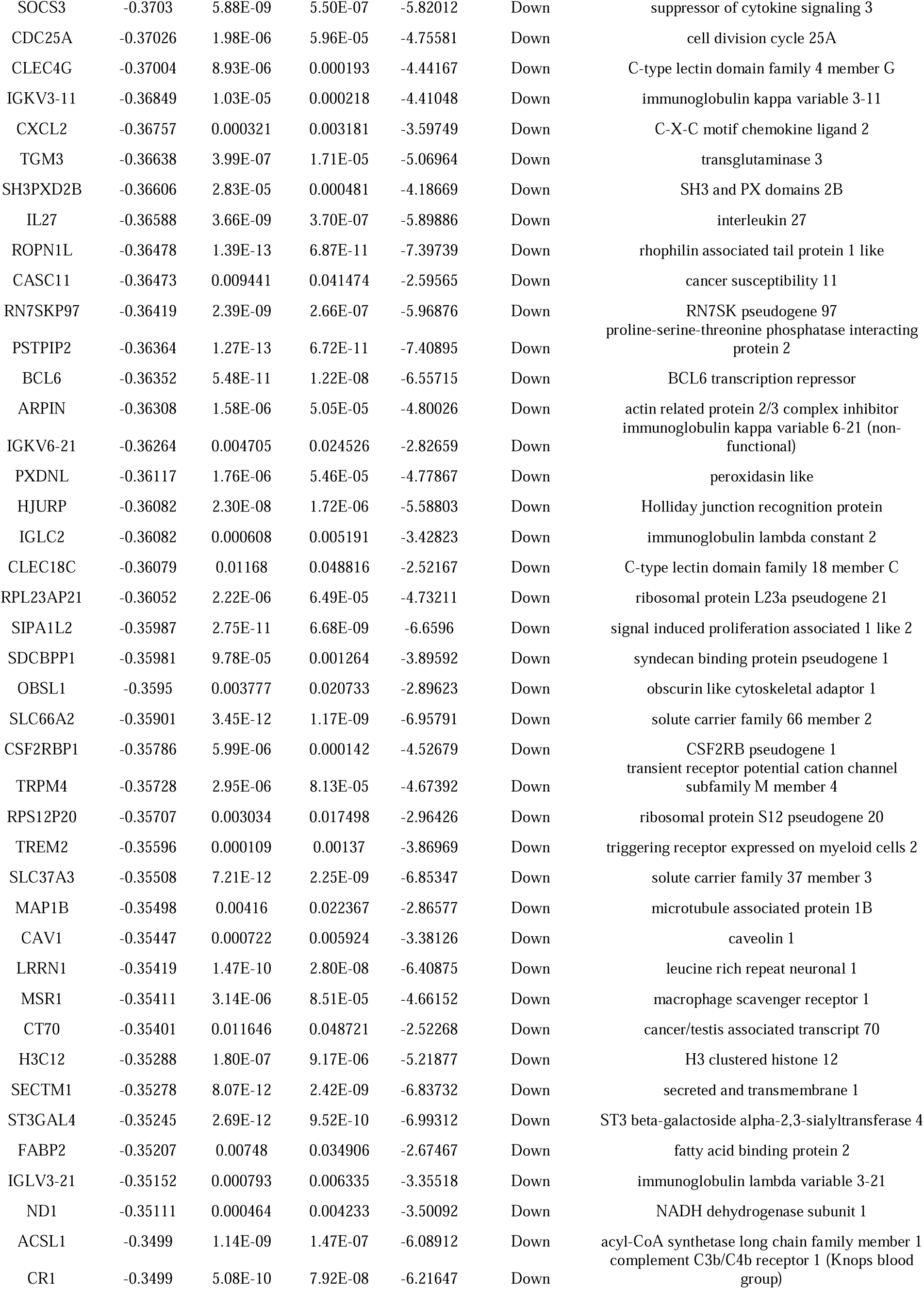
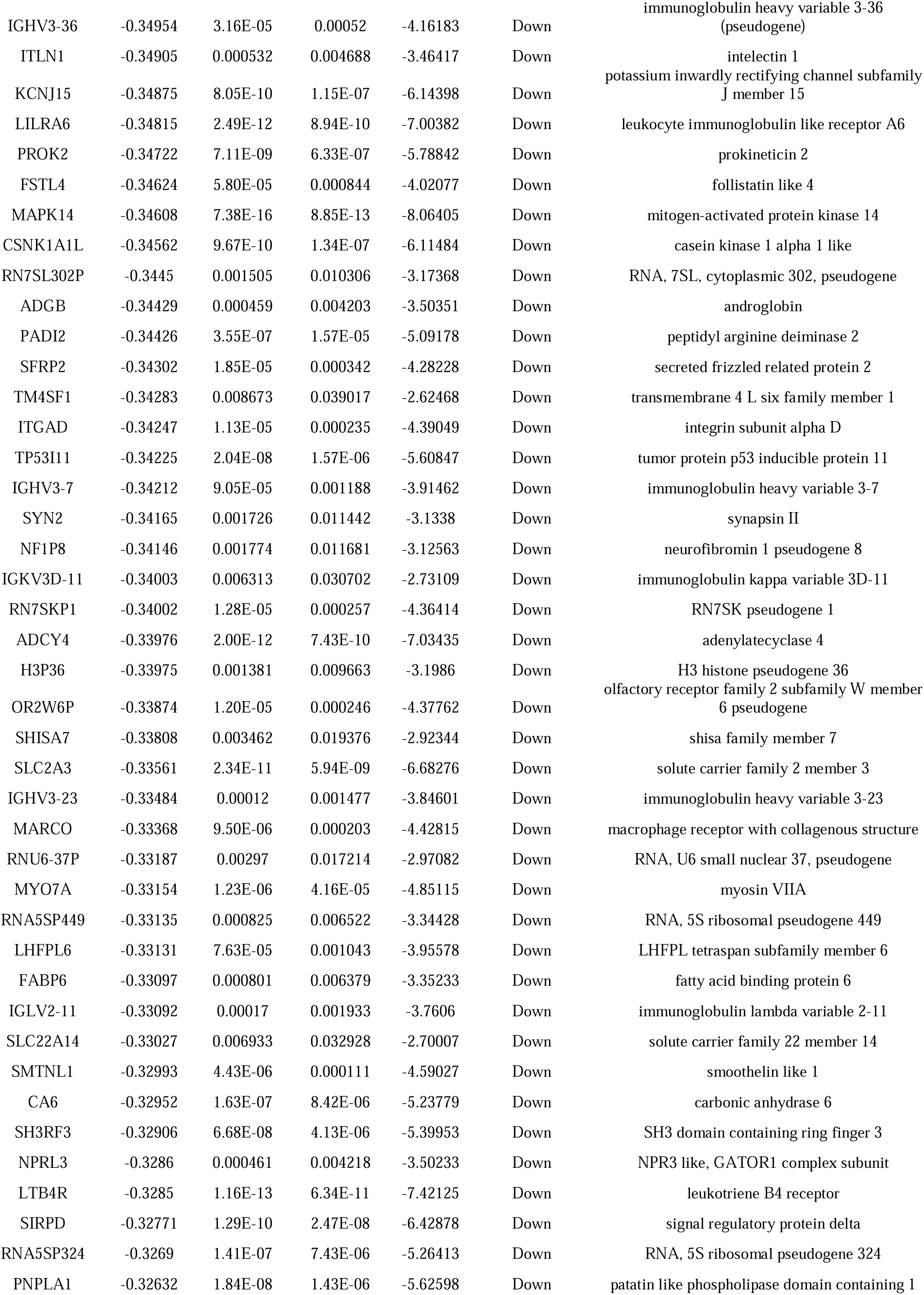
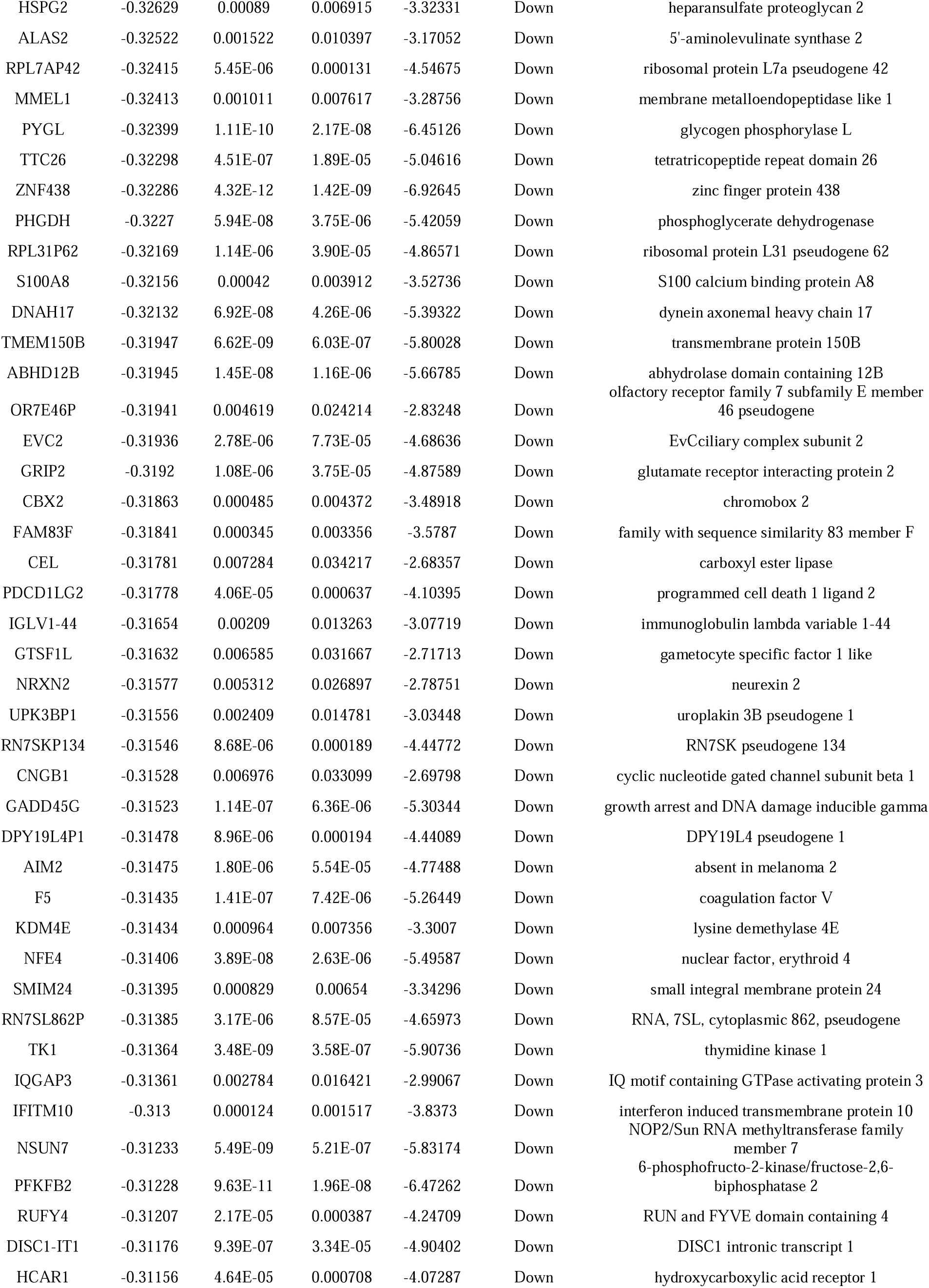
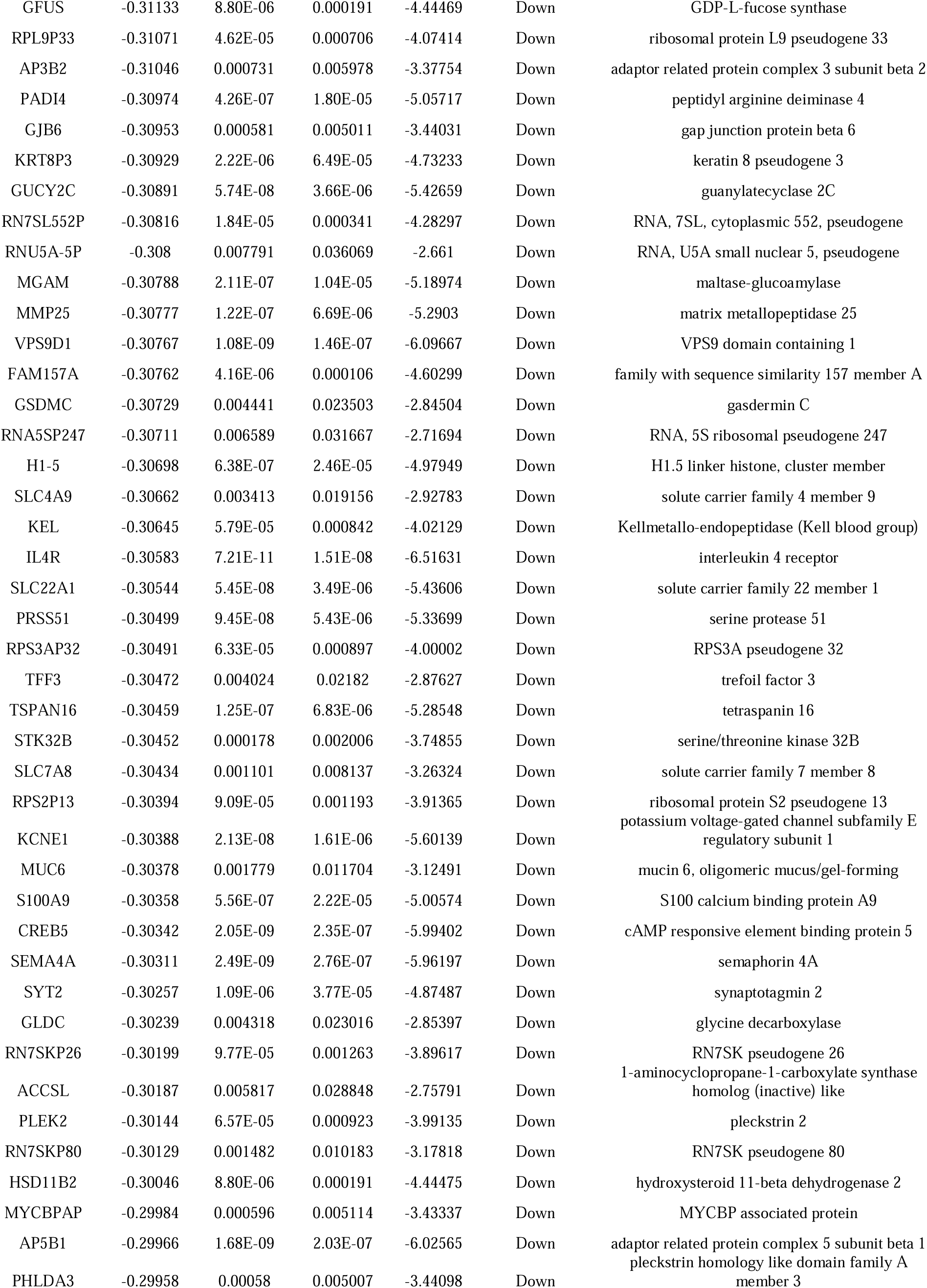
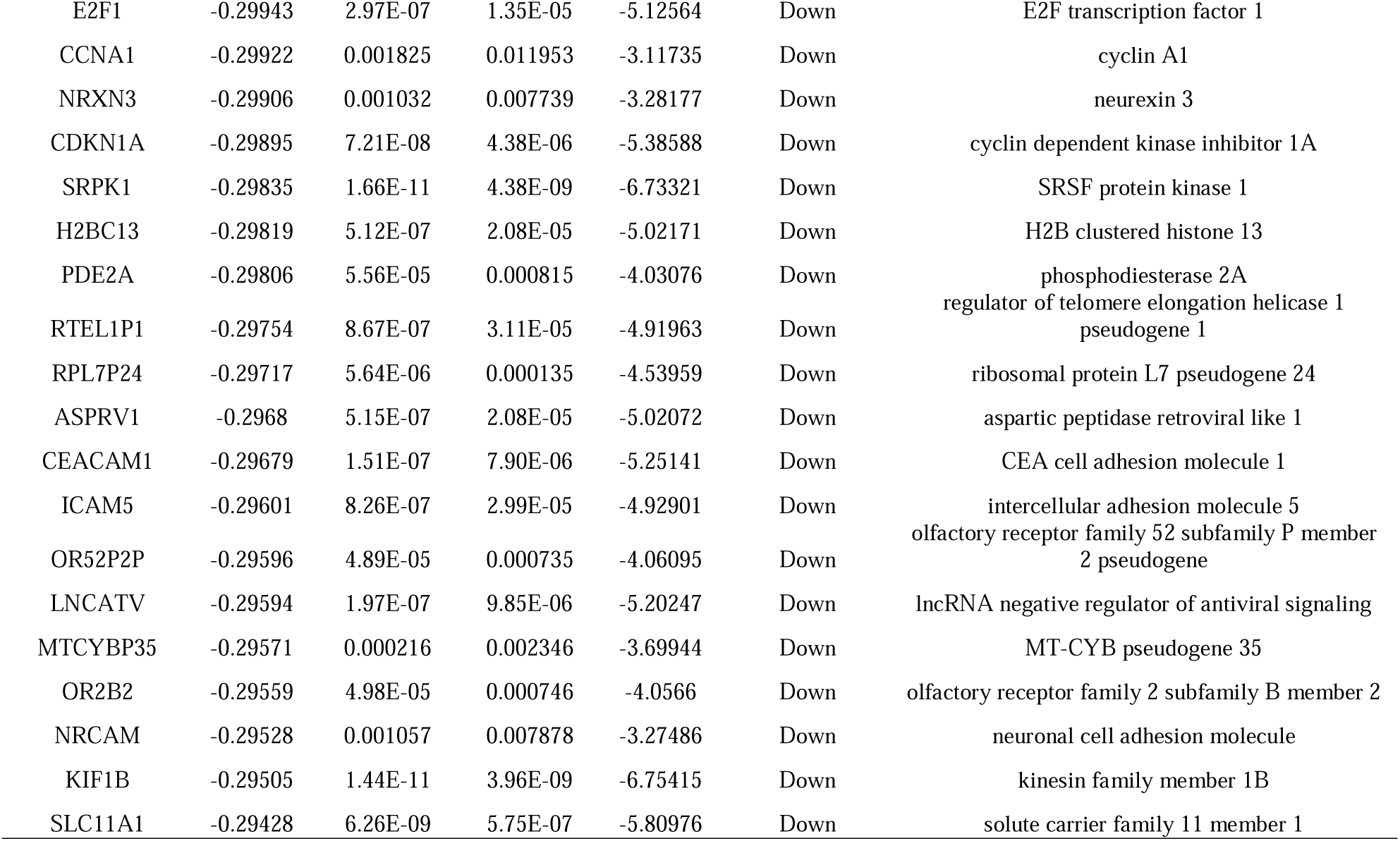
The statistical metrics for key differentially expressed genes (DEGs)

**Fig. 1.**
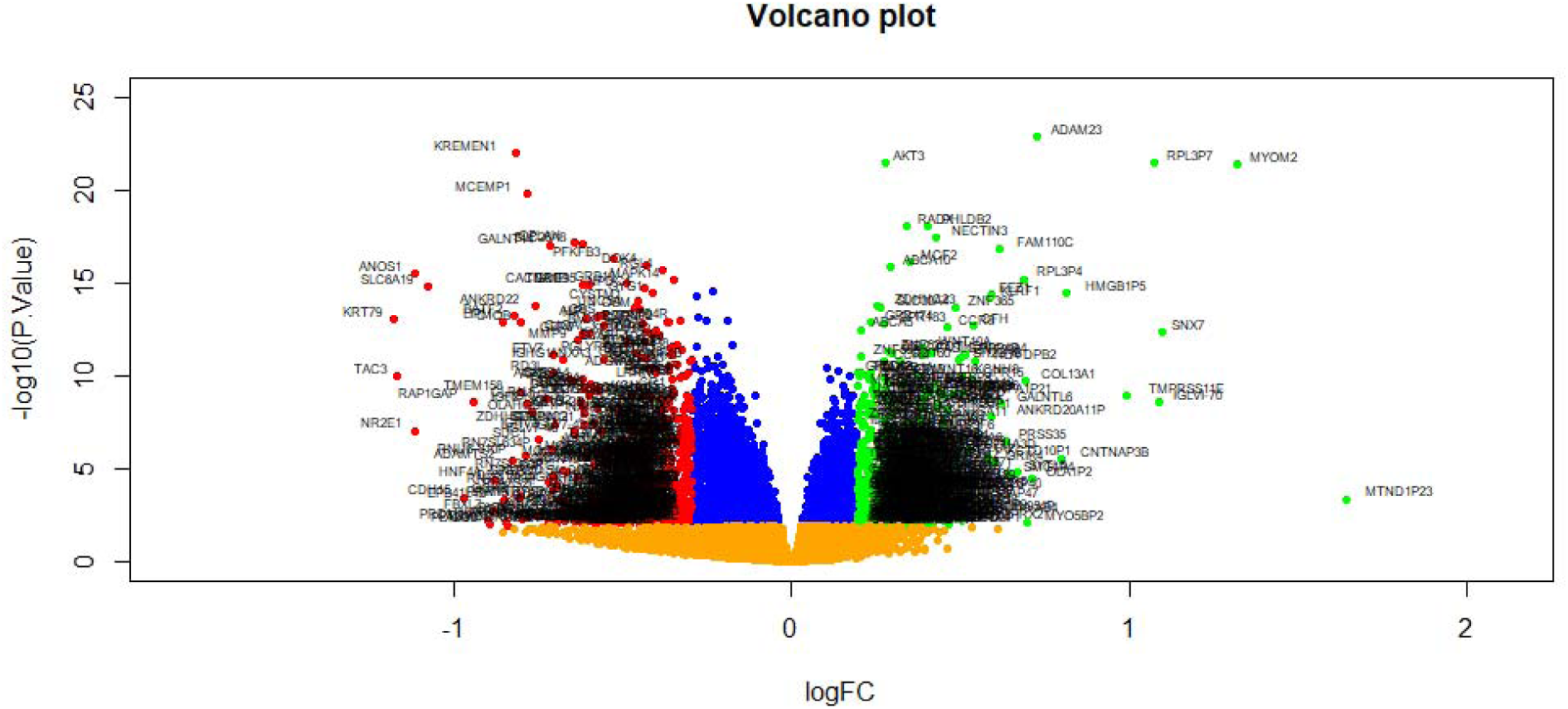
Volcano plot of differentially expressed genes. Genes with a significant change of more than two-fold were selected. Green dot represented up regulated significant genes and red dot represented down regulated significant genes.

**Fig. 2.**
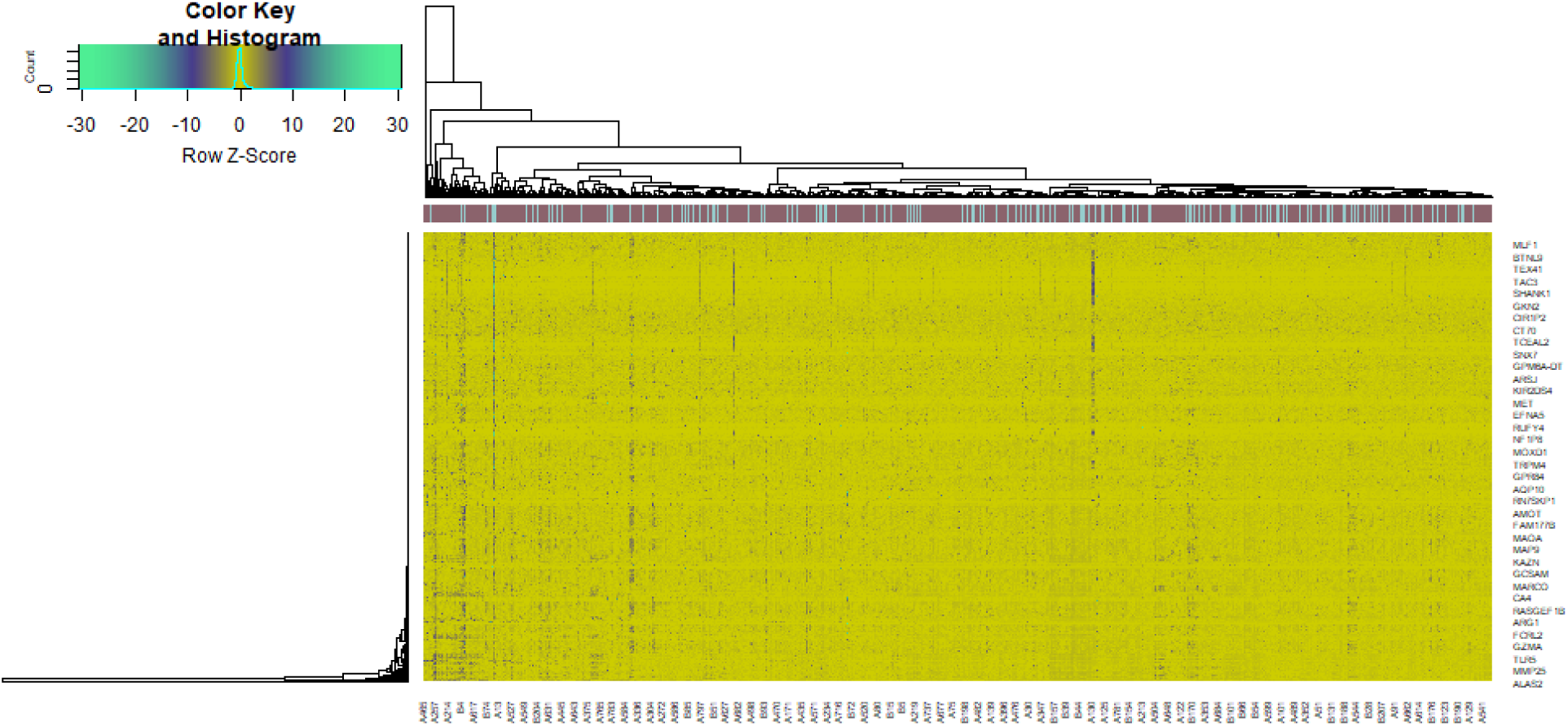
Heat map of differentially expressed genes. Legend on the top left indicate log fold change of genes. (A1 – A807 = IBD samples; B1 – B209 = normal control samples)

### GO and pathway enrichment analyses of DEGs

To determine the biological features of DEGs, GO and pathway enrichment analysis was accomplished by g:Profiler online tools. The BP analysis revealed that the DEGs were major enriched in multicellular organismal process, regulation of cellular process, response to stimulus and developmental process (Table 2). The CC analysis showed that DEGs were enriched in cell periphery, membrane, plasma membrane and cellular anatomical entity (Table 2). Changes in MF of DEGs were significantly enriched in molecular transducer activity, signaling receptor binding and identical protein binding (Table 2). The results of REACTOME pathway enrichment analysis revealed that DEGs were mainly involved in GPCR ligand binding, class A/1 (Rhodopsin-like receptors), immune system and hemostasis (Table 3).

**Table 2.**
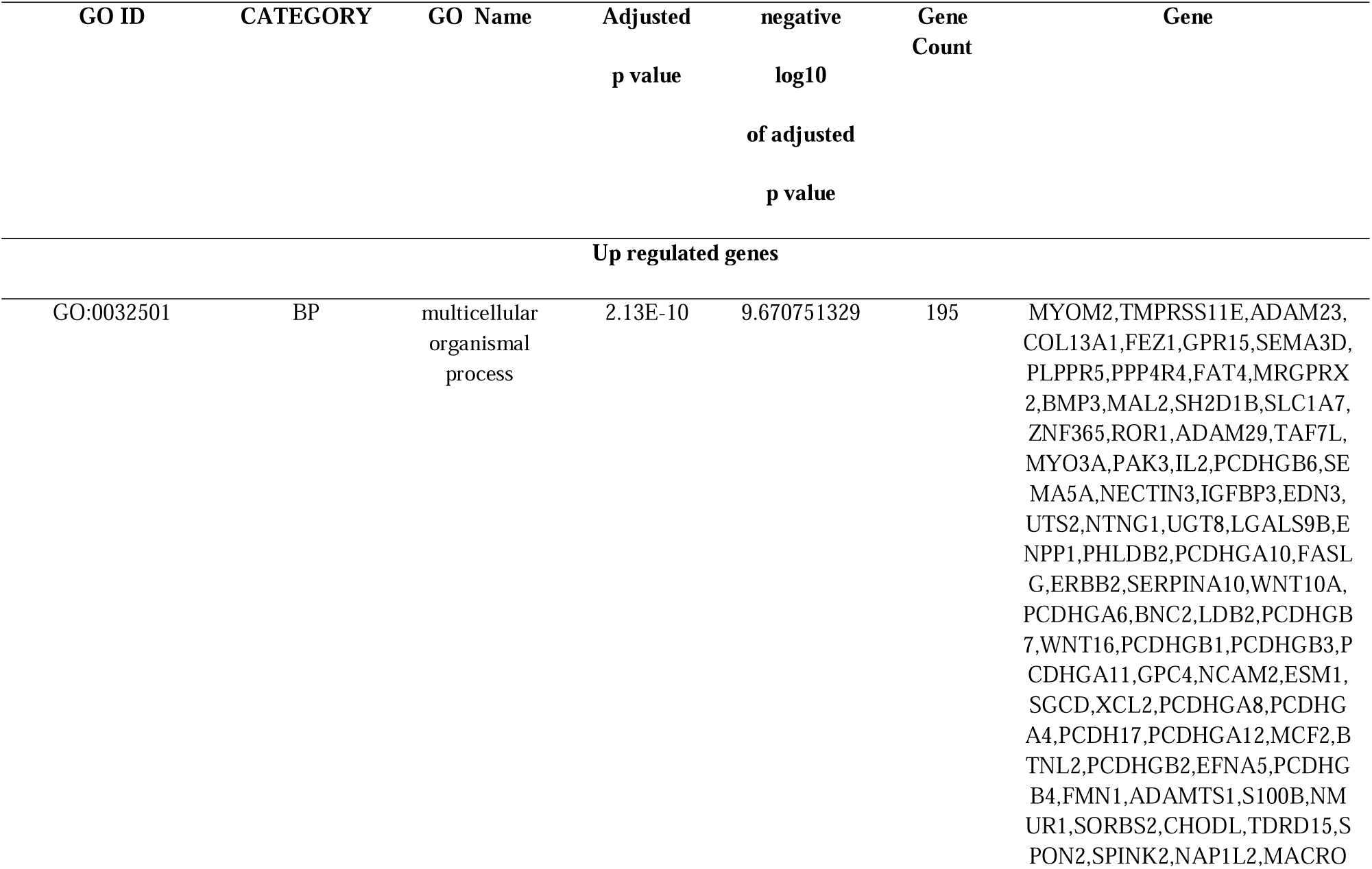
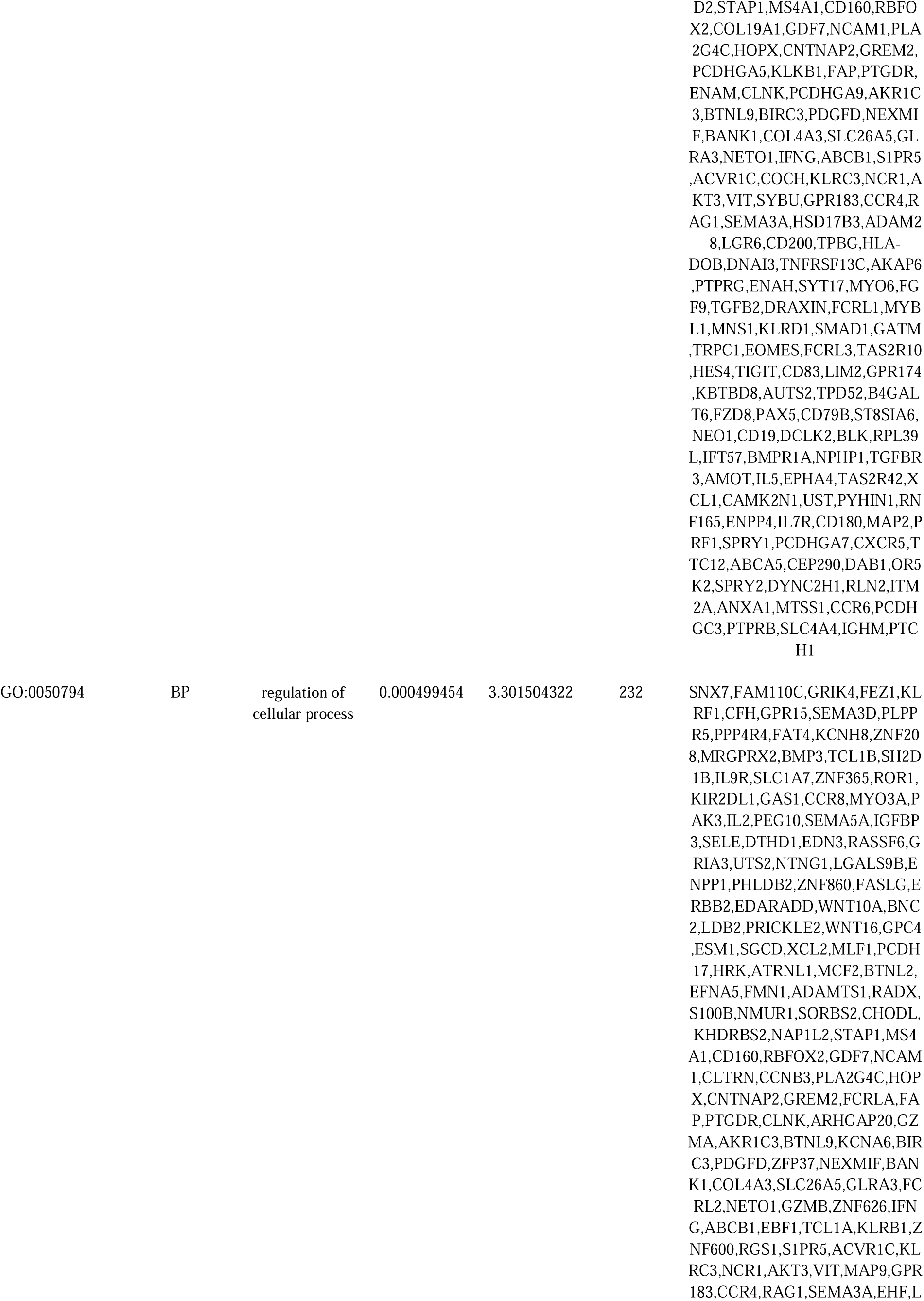
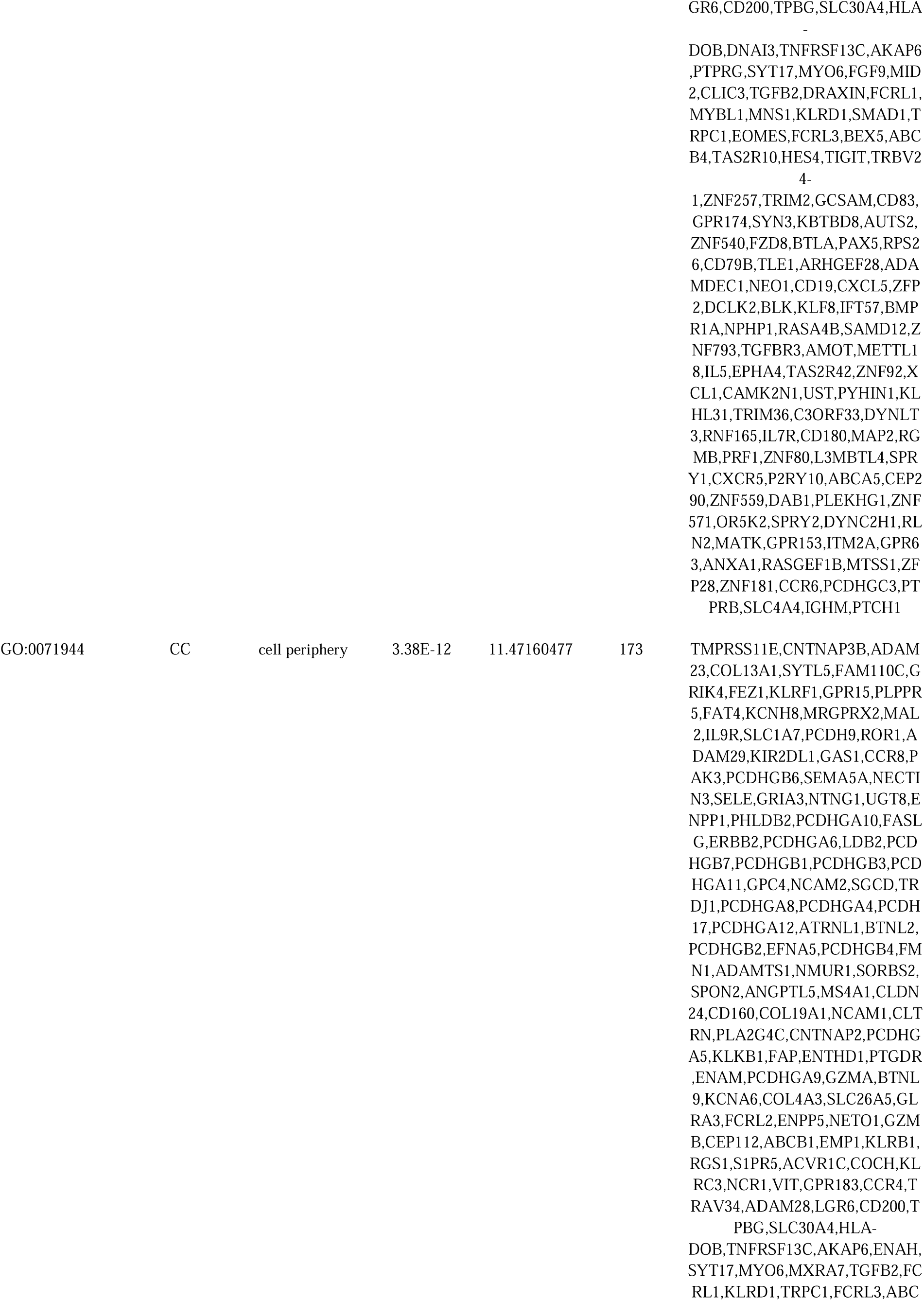
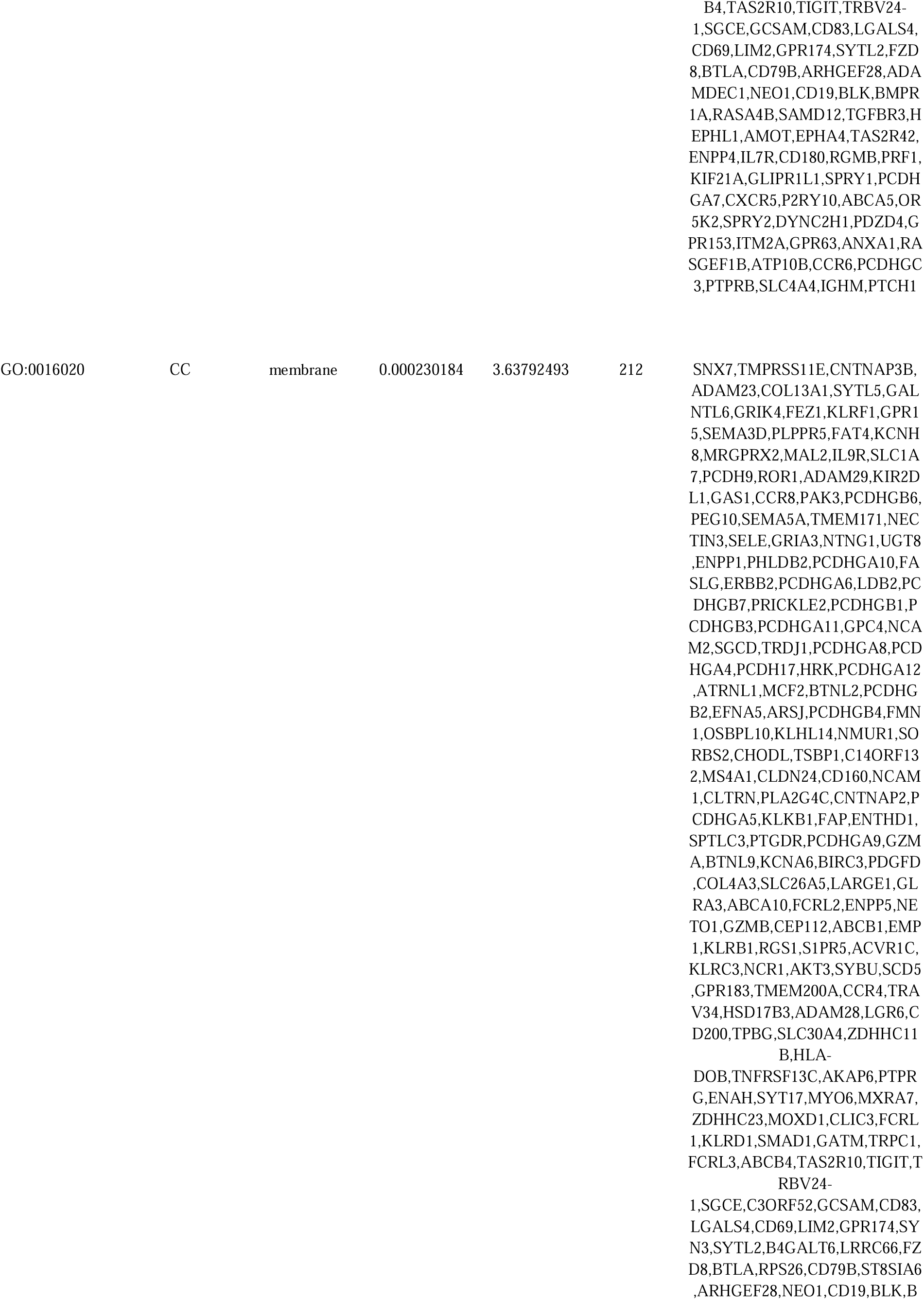
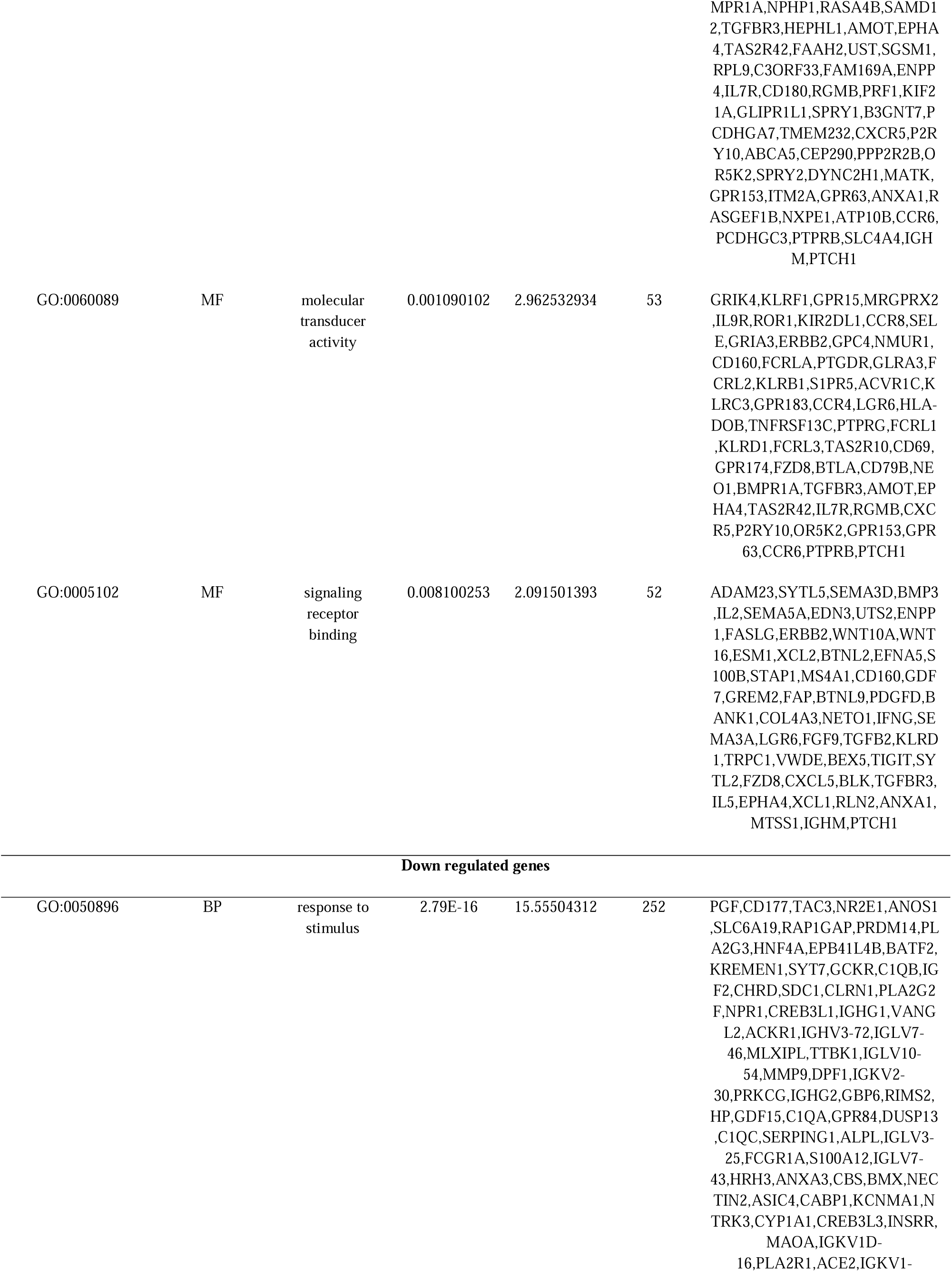
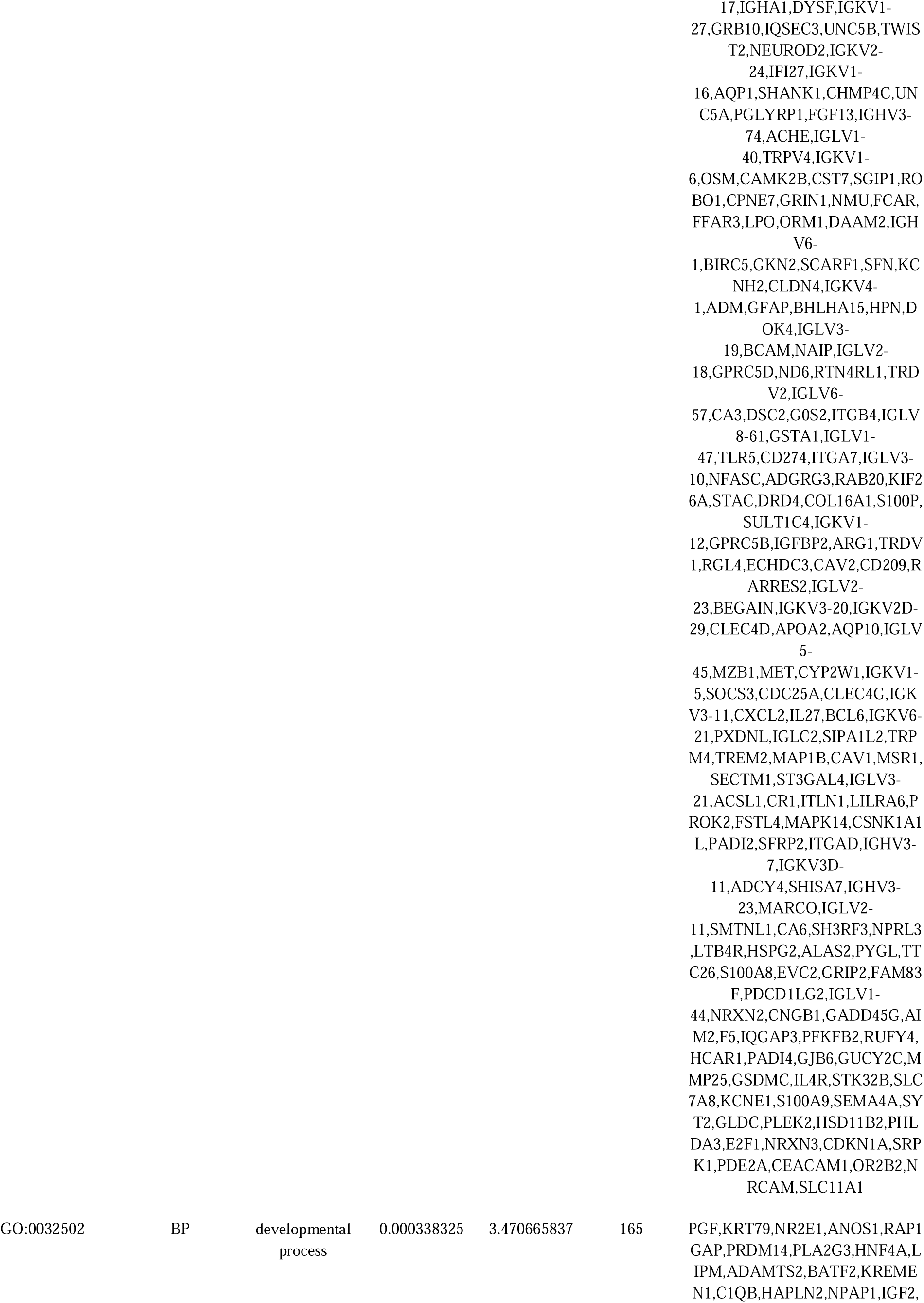
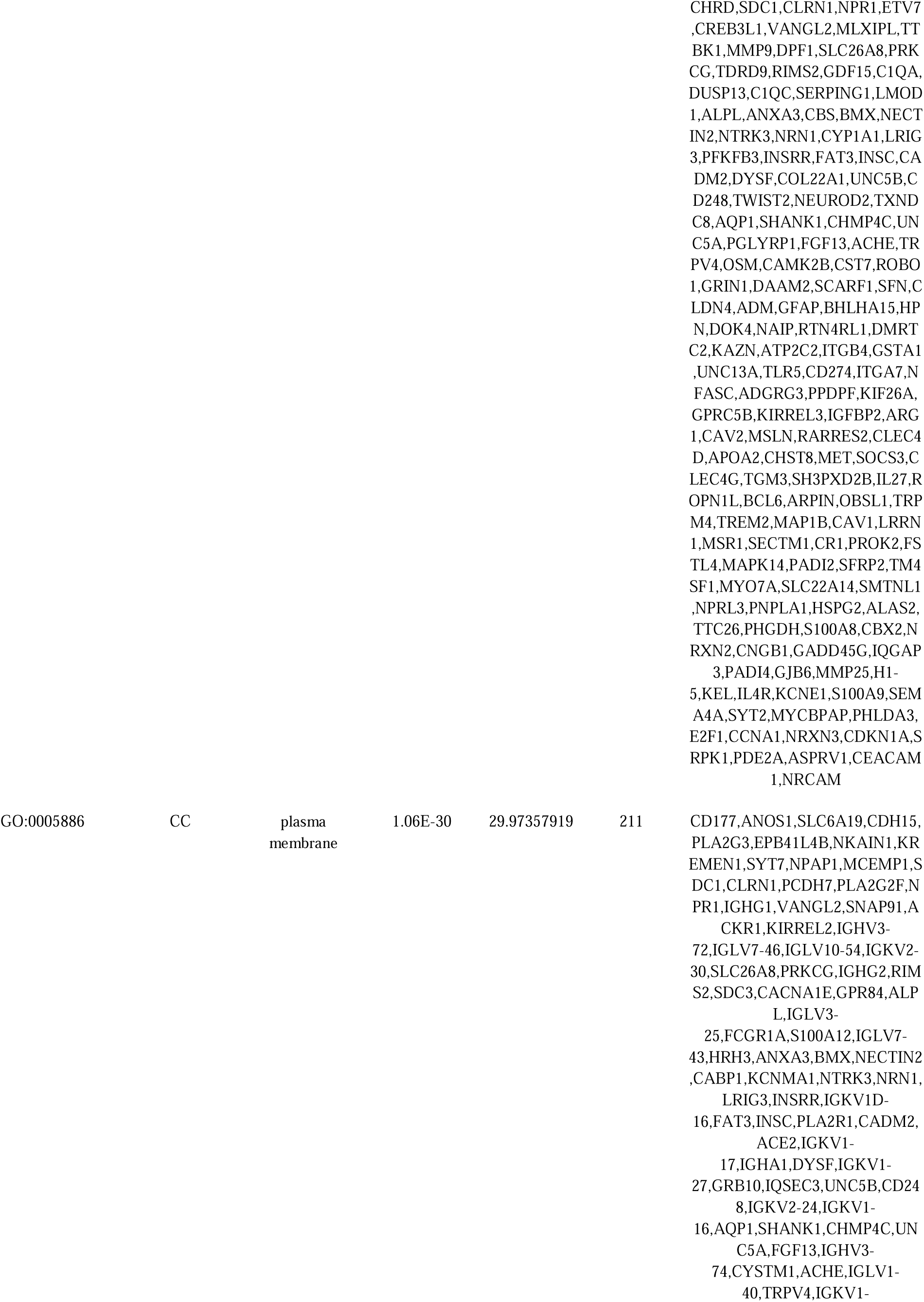
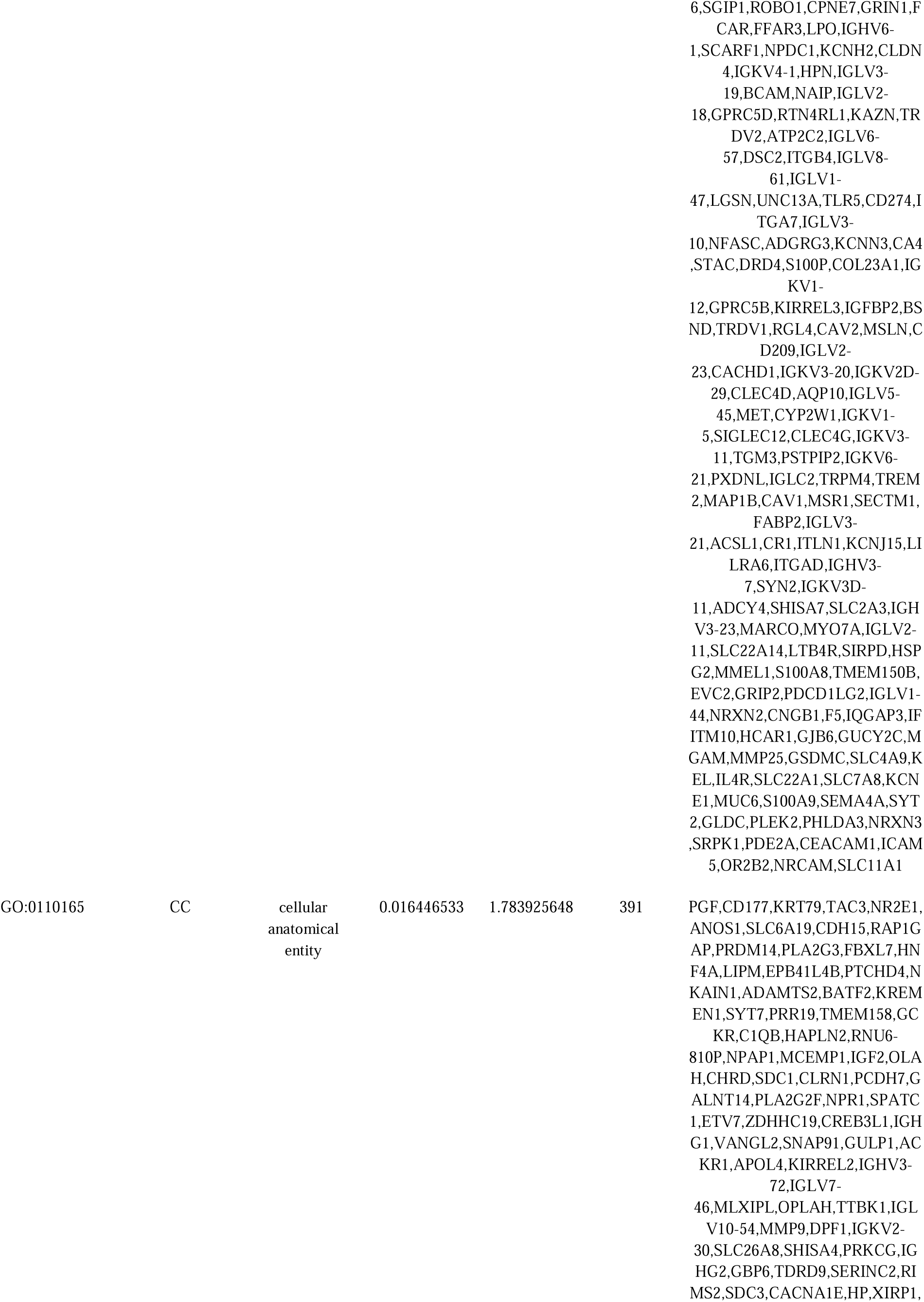
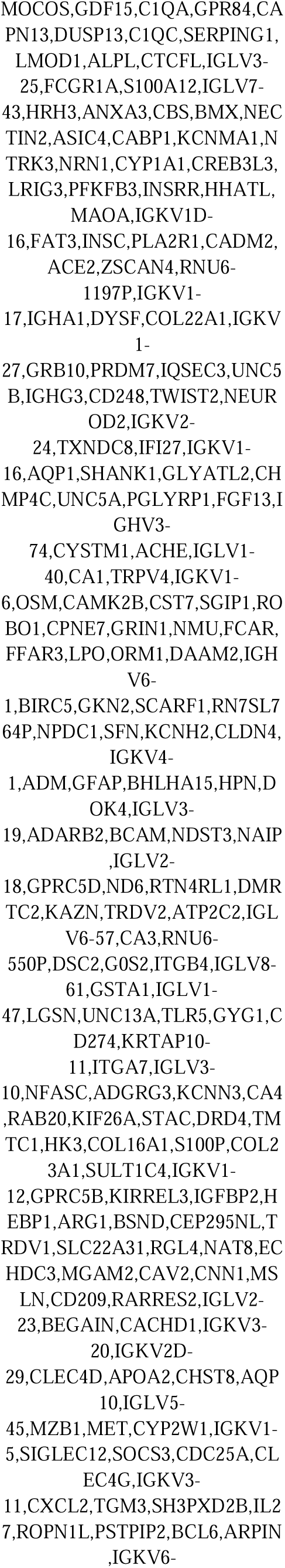
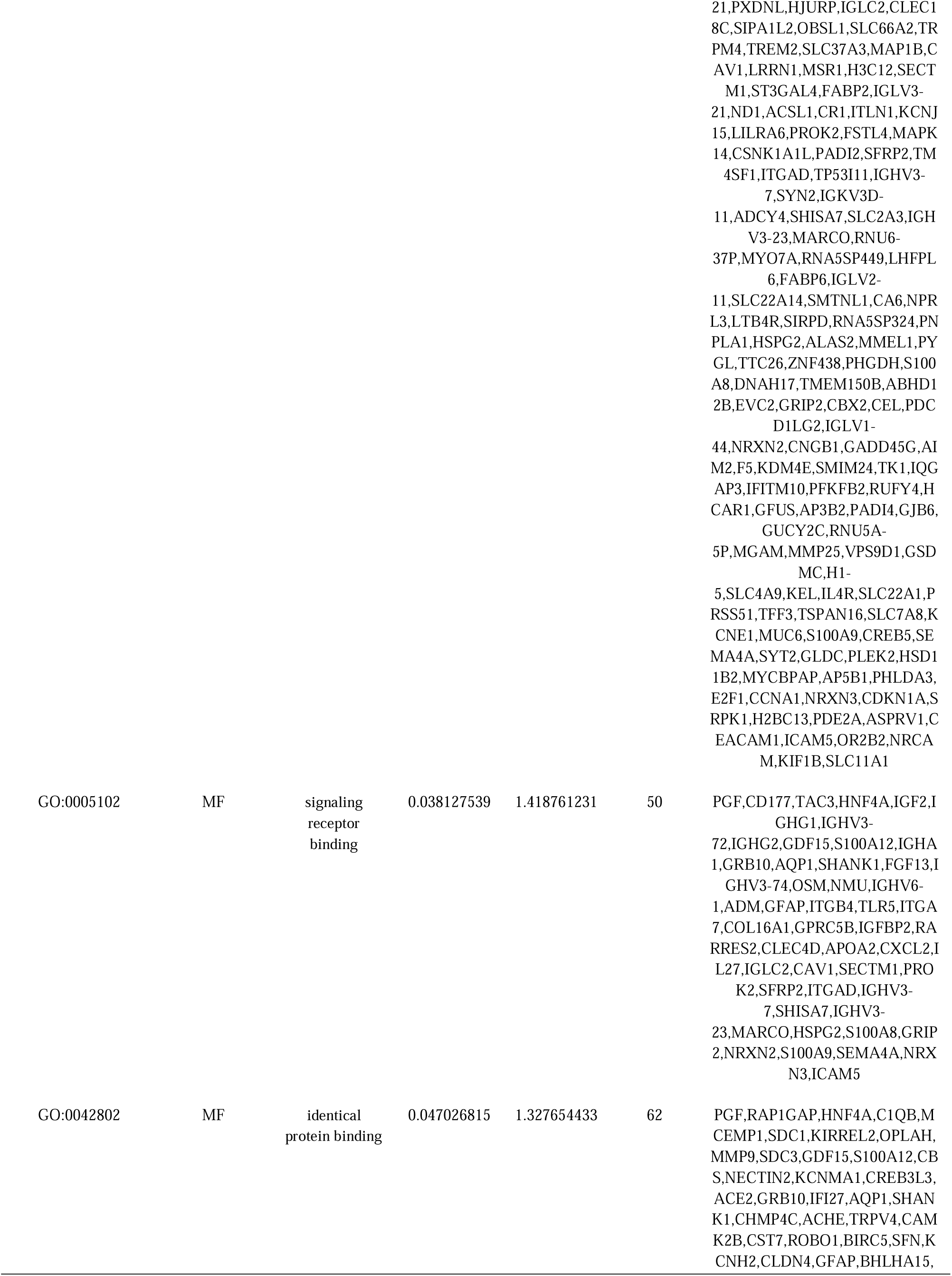
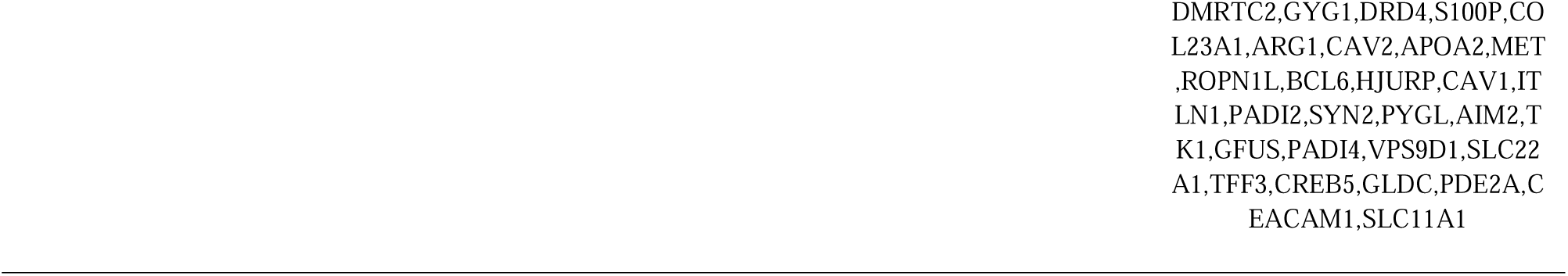
The enriched GO terms of the up and down regulated differentially expressed genes

**Table 3.**
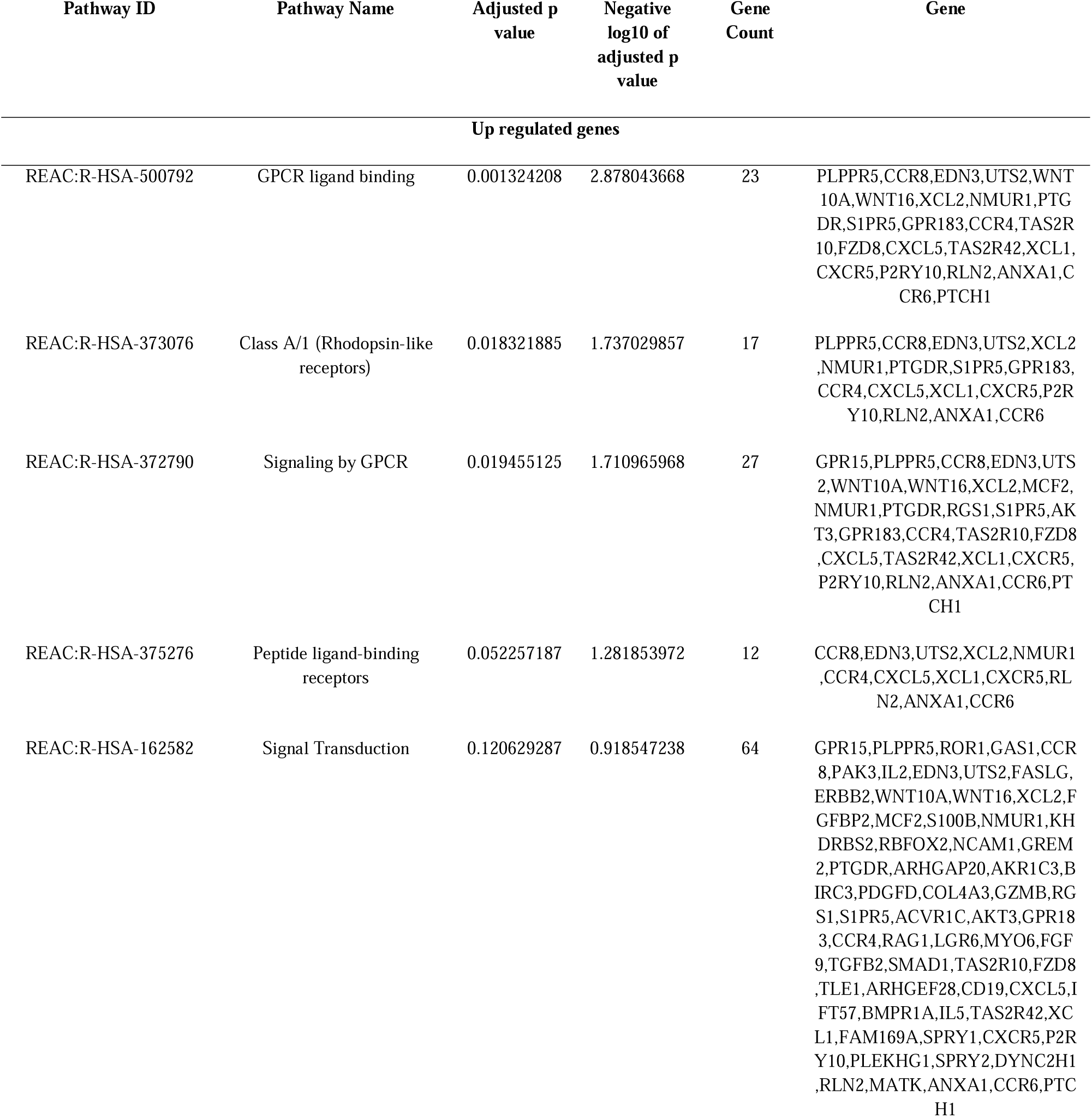
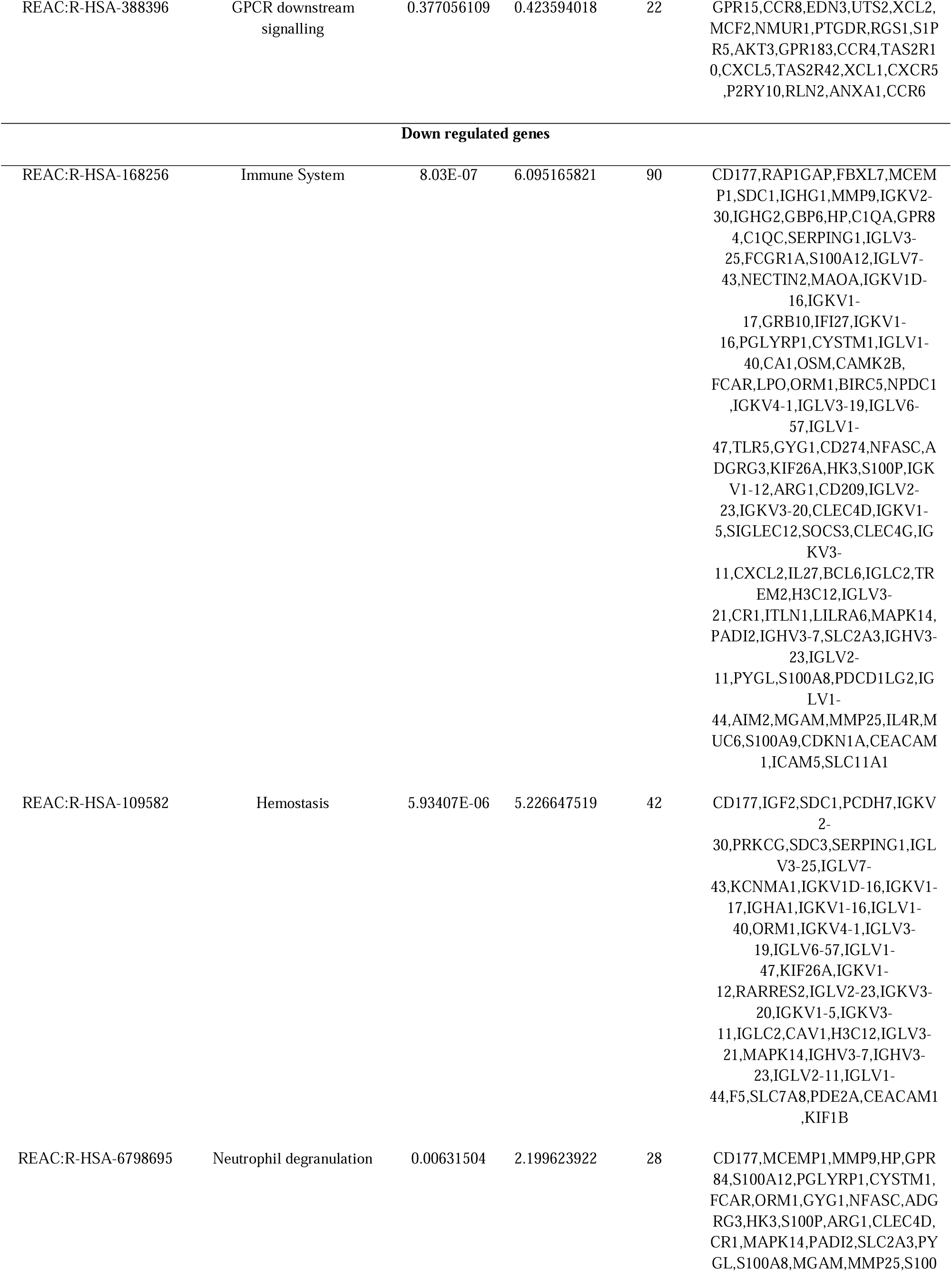
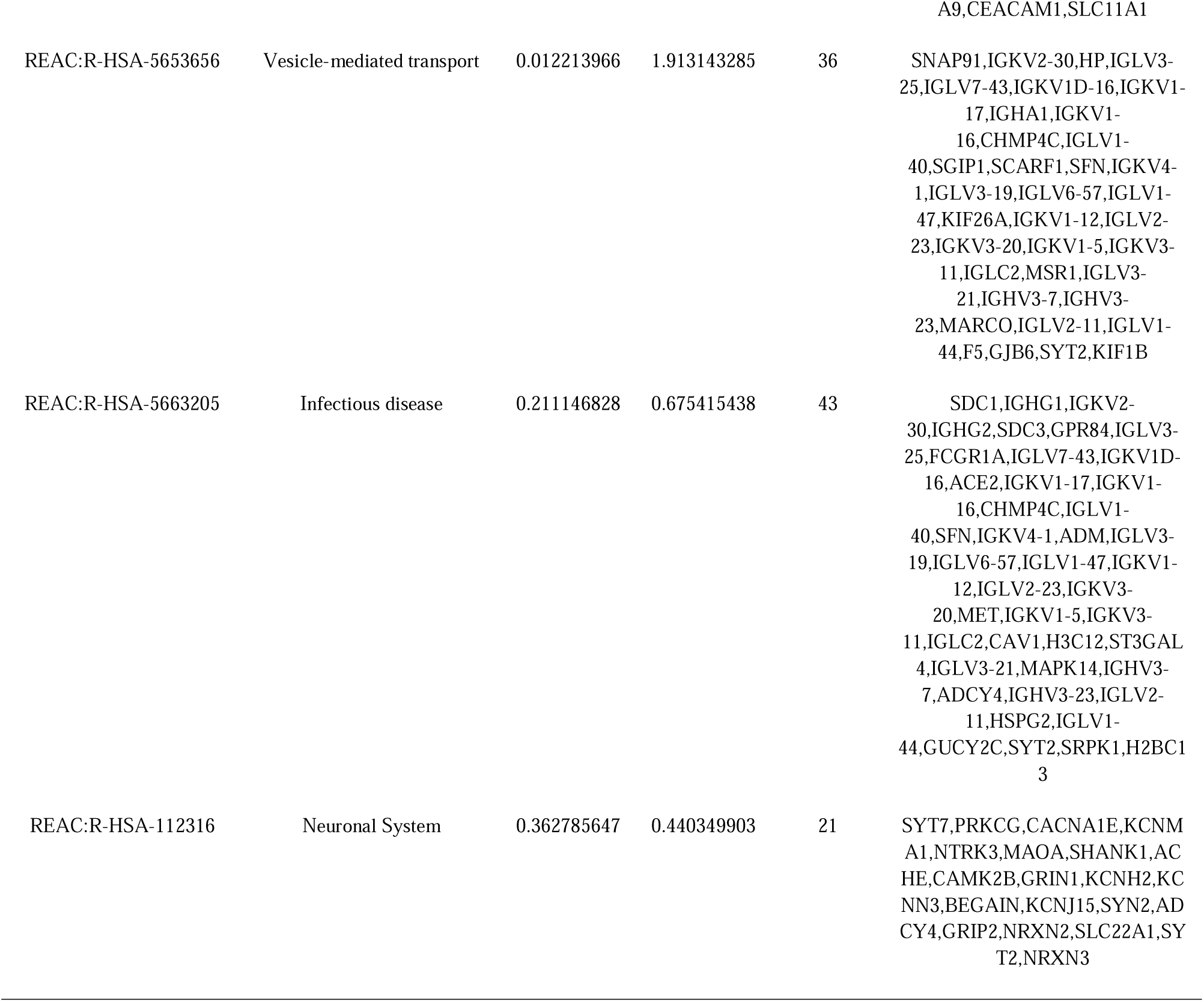
The enriched pathway terms of the up and down regulated differentially expressed genes

### Construction of the PPI network and module analysis

Uploading 957 DEGs into the IMEX interactome database, a PPI network with the desired interaction scoreL>L0.48 was constructed and the network containing 4783 nodes and 8031 edges was visualized via Cytoscape software (Fig. 3). Then the most remarkable hub genes were recognized by Network Analyzer plug-in according to the maximal degree, betweenness, stress and closeness topology analysis methods. The top nodes appraised as hub genes: IL7R, ERBB2, SMAD1, RPS26, TLE1, HNF4A, CDKN1A, SRPK1, H3C12 and SFN (Table 4). The key modules were obtained using PEWCC. The two key modules with the highest degree were screened, and the GO and pathway enrichment of genes in these two modules was analyzed using g:Profiler online tools. Module 1 contained 68 nodes and 133 edges (Fig. 4A), mainly involved in membrane and regulation of cellular process. Module 2 was comprised of 21 nodes and 41 edges (Fig. 4B), which was associated with response to stimulus.

**Fig. 3.**
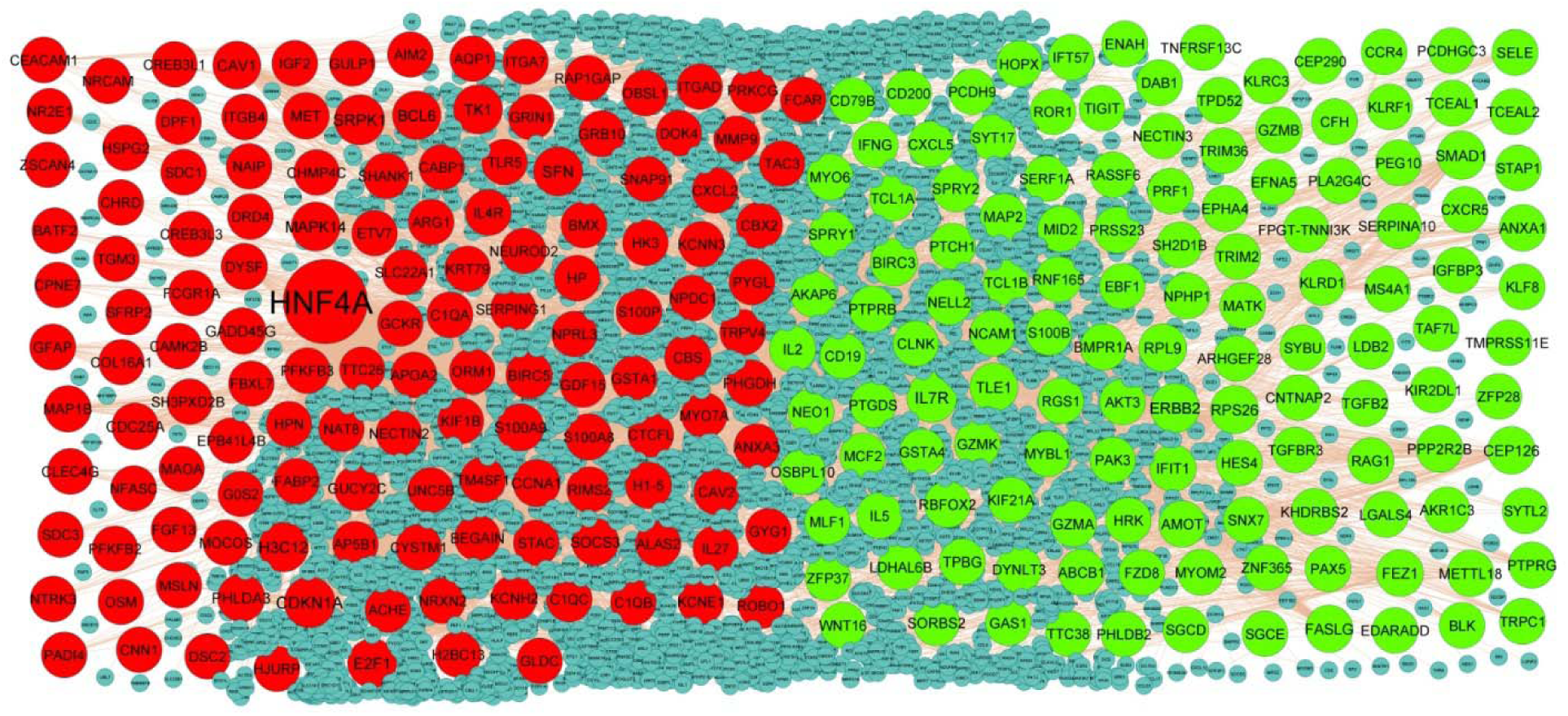
PPI network of DEGs. Up regulated genes are marked in parrot green; down regulated genes are marked in red

**Table 4.**
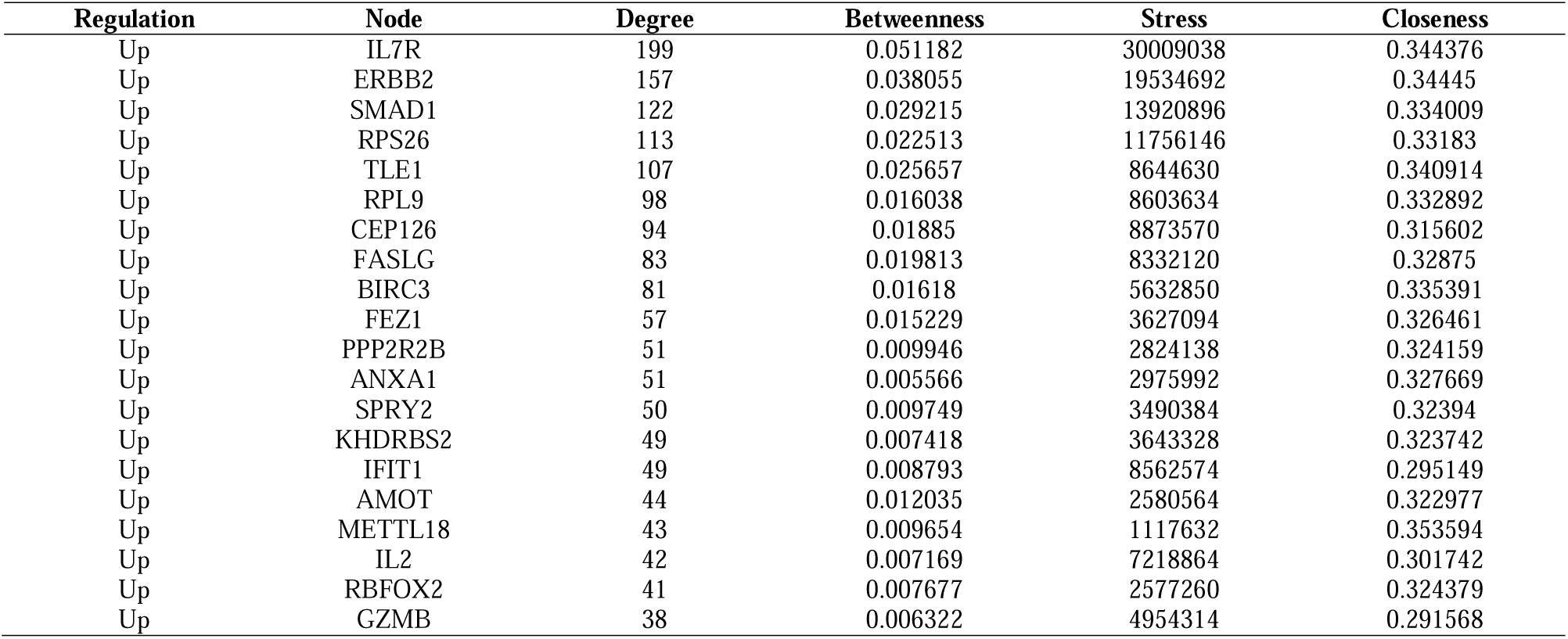
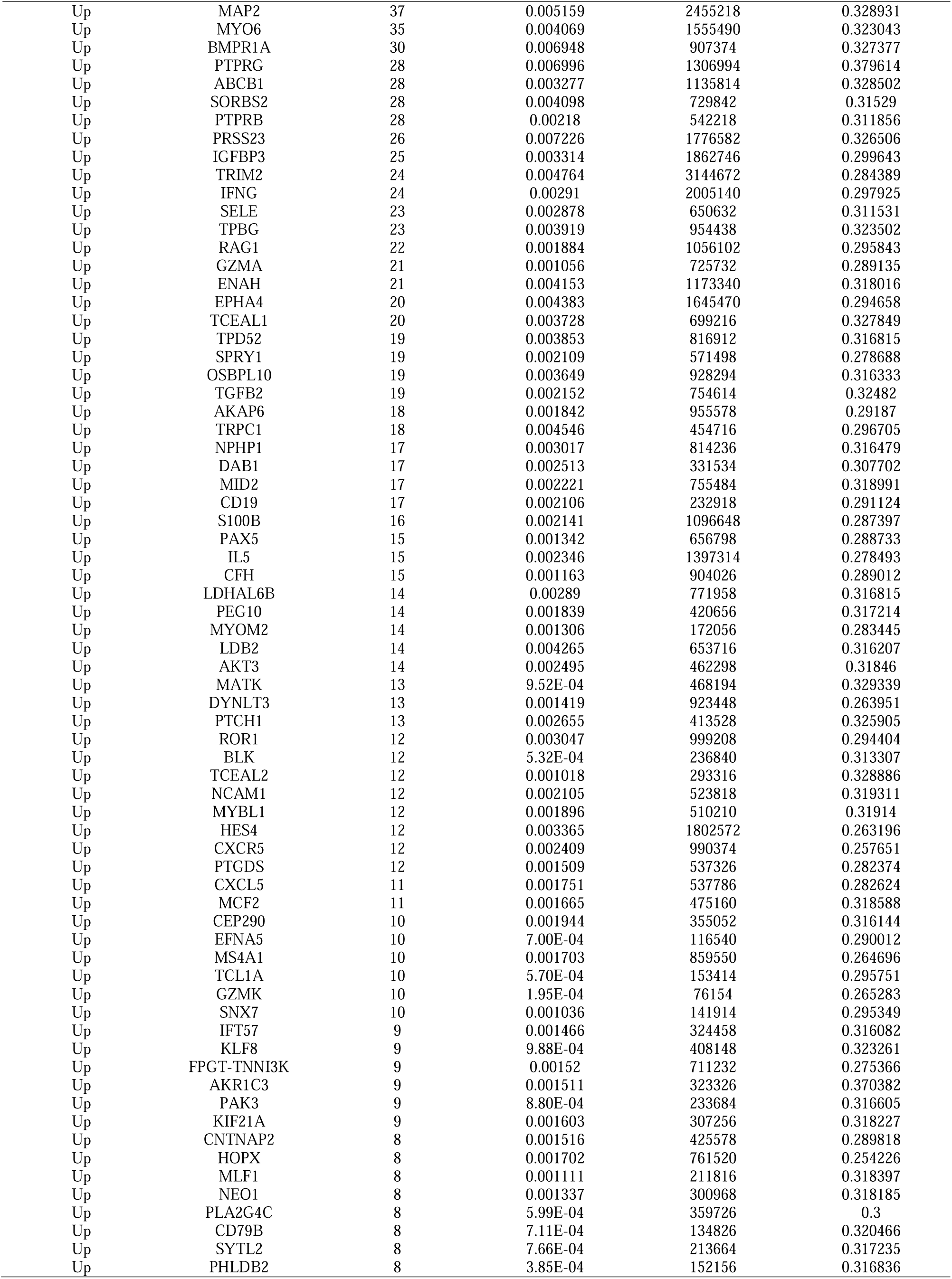
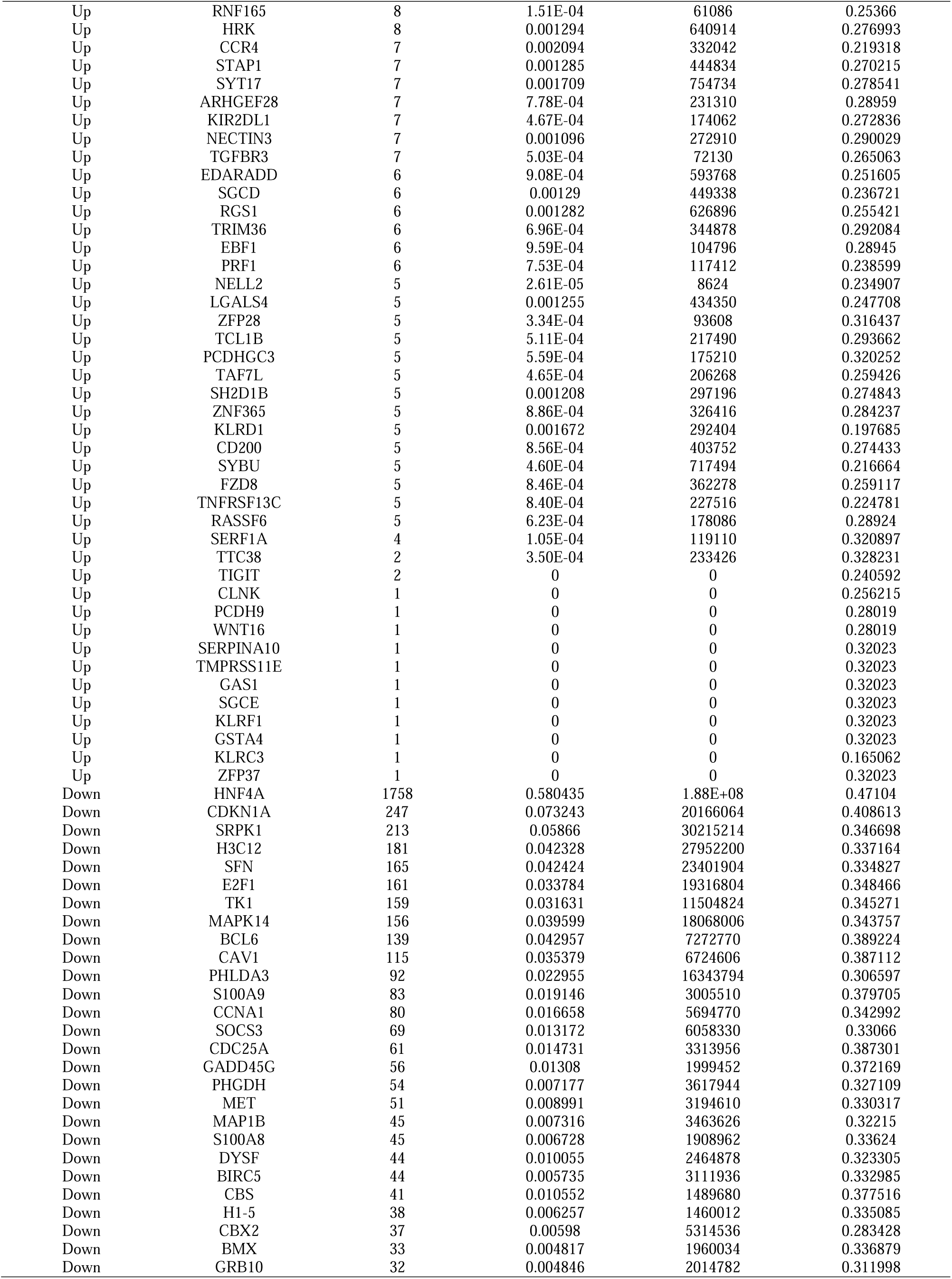
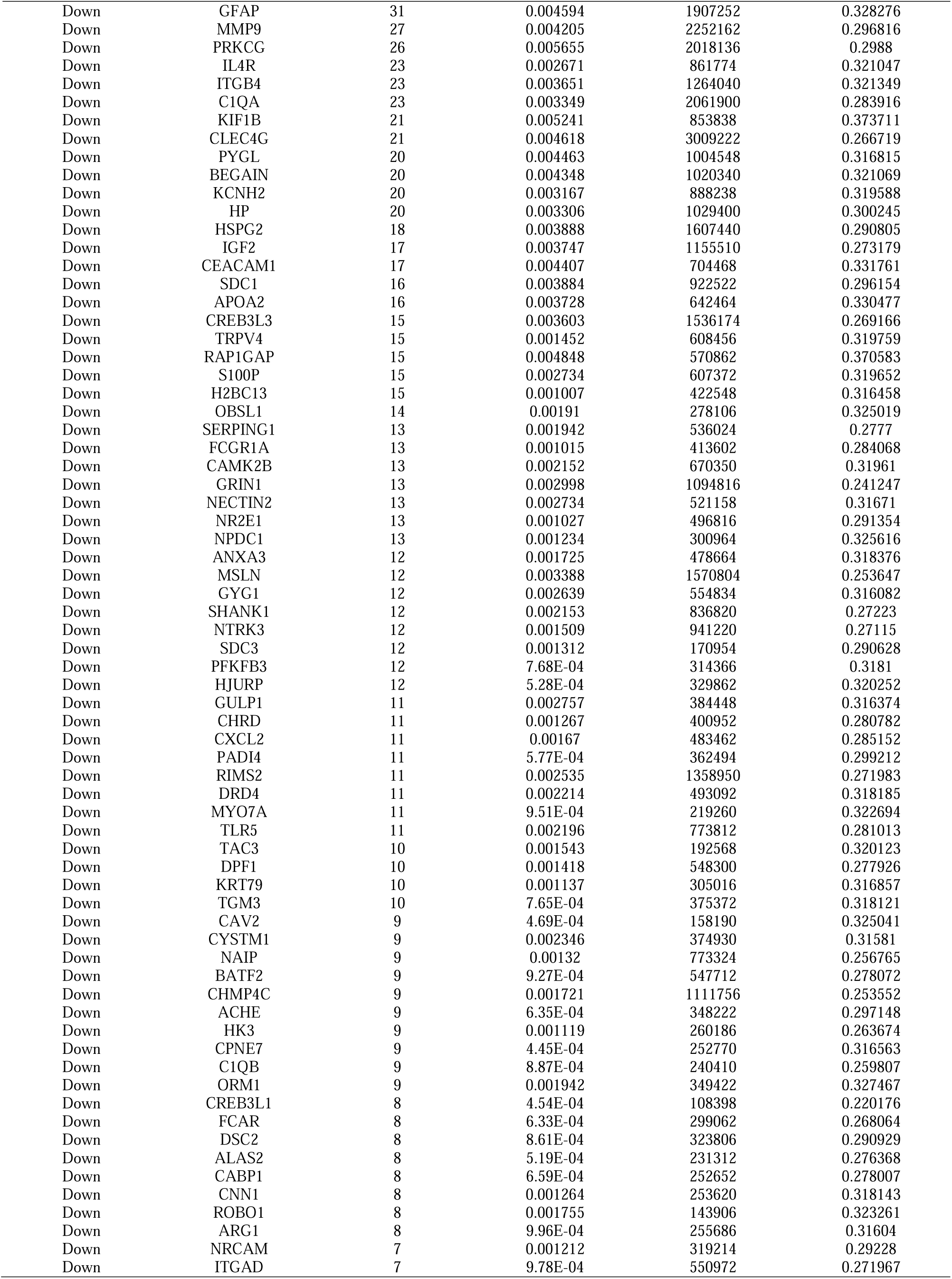
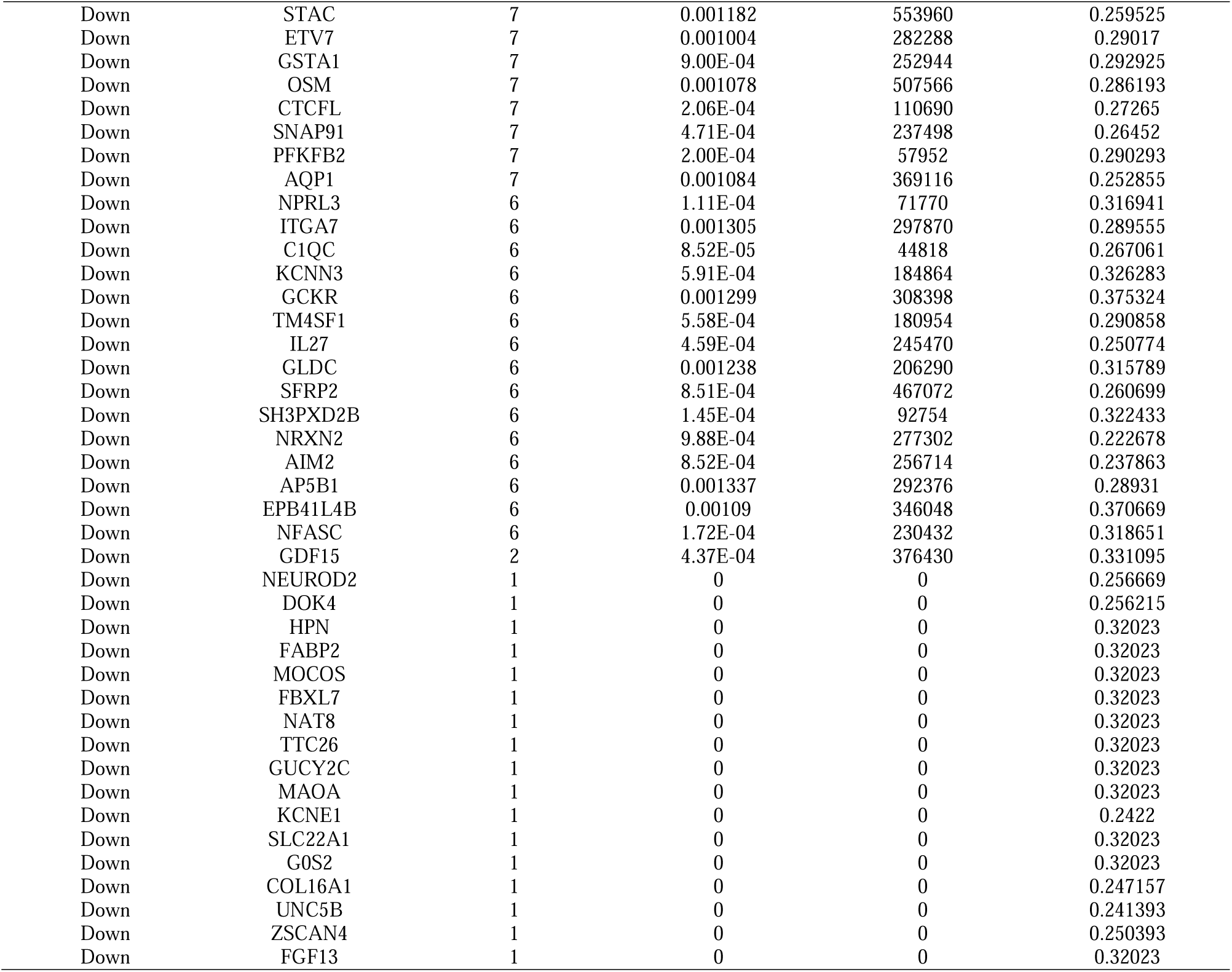
Topology table for up and down regulated genes

**Fig. 4.**
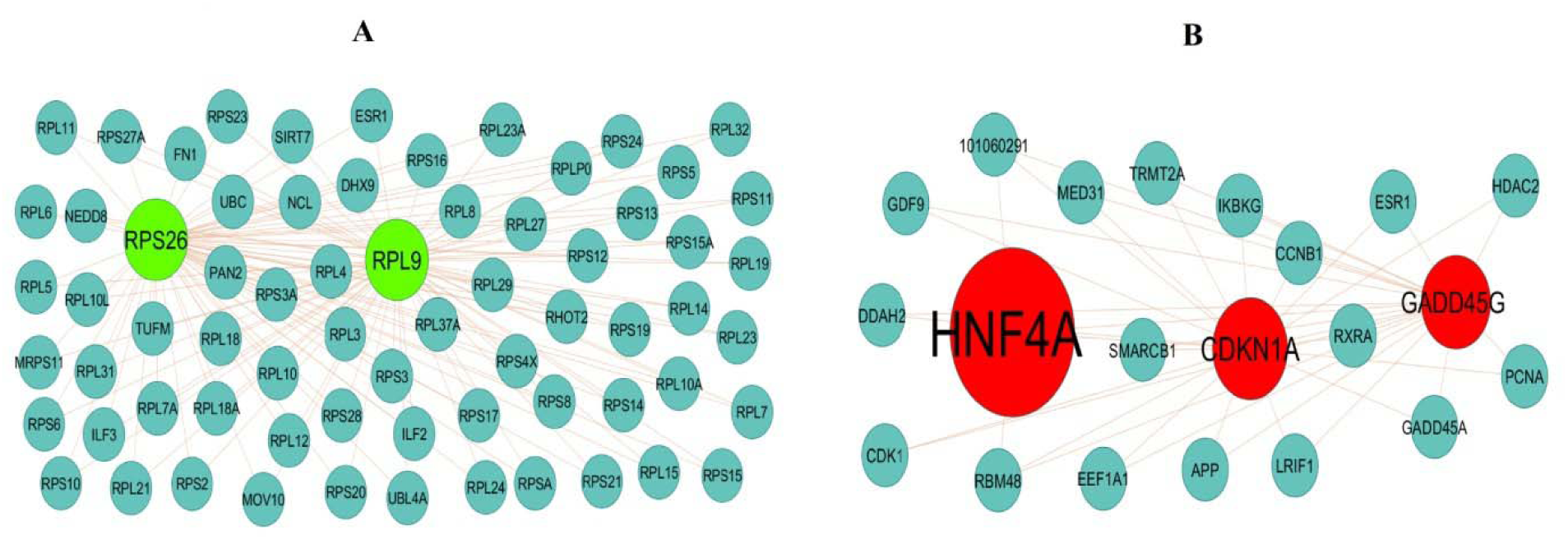
Modules selected from the DEG PPI between patients with FSGS and normal controls. (A) The most significant module was obtained from PPI network with 68 nodes and 133 edges for up regulated genes (B) The most significant module was obtained from PPI network with 21 nodes and 41 edges for down regulated genes. Up regulated genes are marked in parrot green; down regulated genes are marked in red

### Construction of the miRNA-hub gene regulatory network

The miRNA-hub gene regulatory networks of the hub genes in IDB were constructed through miRNet database, and 11005 edges involving 2326 nodes (miRNAs: 2061; Hub Genes: 265) were formed in the network (Fig. 5). ERBB2 was regulated by 101 different miRNAs (ex: hsa-mir-3921); BIRC3 was regulated by 83 different miRNAs (ex: hsa-mir-449a); IFIT1 was regulated by 67 different miRNAs (ex: hsa-mir-375); SPRY2 was regulated by 46 different miRNAs (ex: hsa-mir-21-5p); IL7R was regulated by 40 different miRNAs (ex: hsa-mir-32-3p); CDKN1A was regulated by 444 different miRNAs (ex: hsa-mir-5685); PHLDA3 was regulated by 155 different miRNAs (ex: hsa-mir-5693); MAPK14 was regulated by 124 different miRNAs (ex: hsa-mir-153-5p); E2F1 was regulated by 121 different miRNAs (ex: hsa-mir-4283); CDC25A was regulated by 121 different miRNAs (ex: hsa-mir-21) (Table 5).

**Fig. 5.**
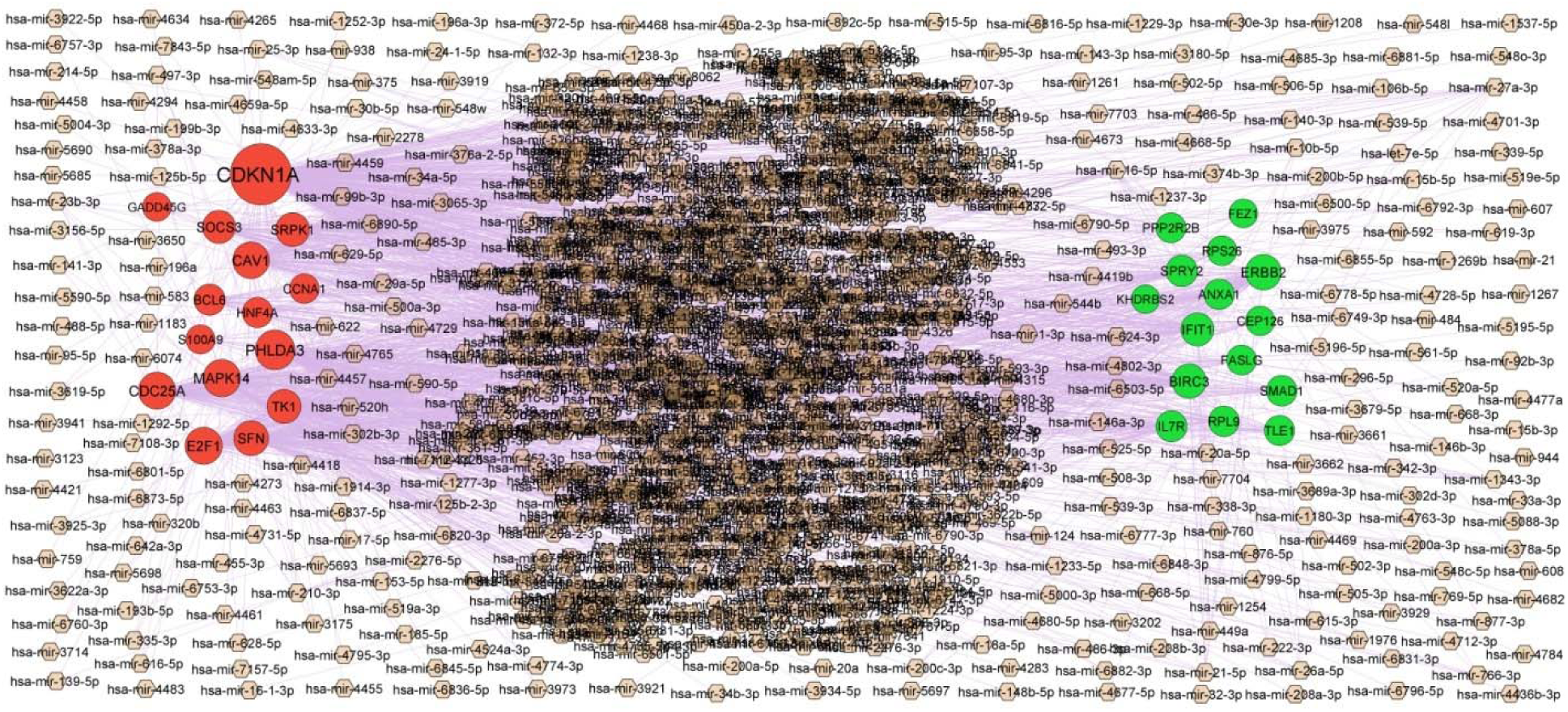
Target gene miRNA regulatory network between target genes. The pink color diamond nodes represent the key miRNAs; up regulated genes are marked in green; down regulated genes are marked in red.

**Table 5.**
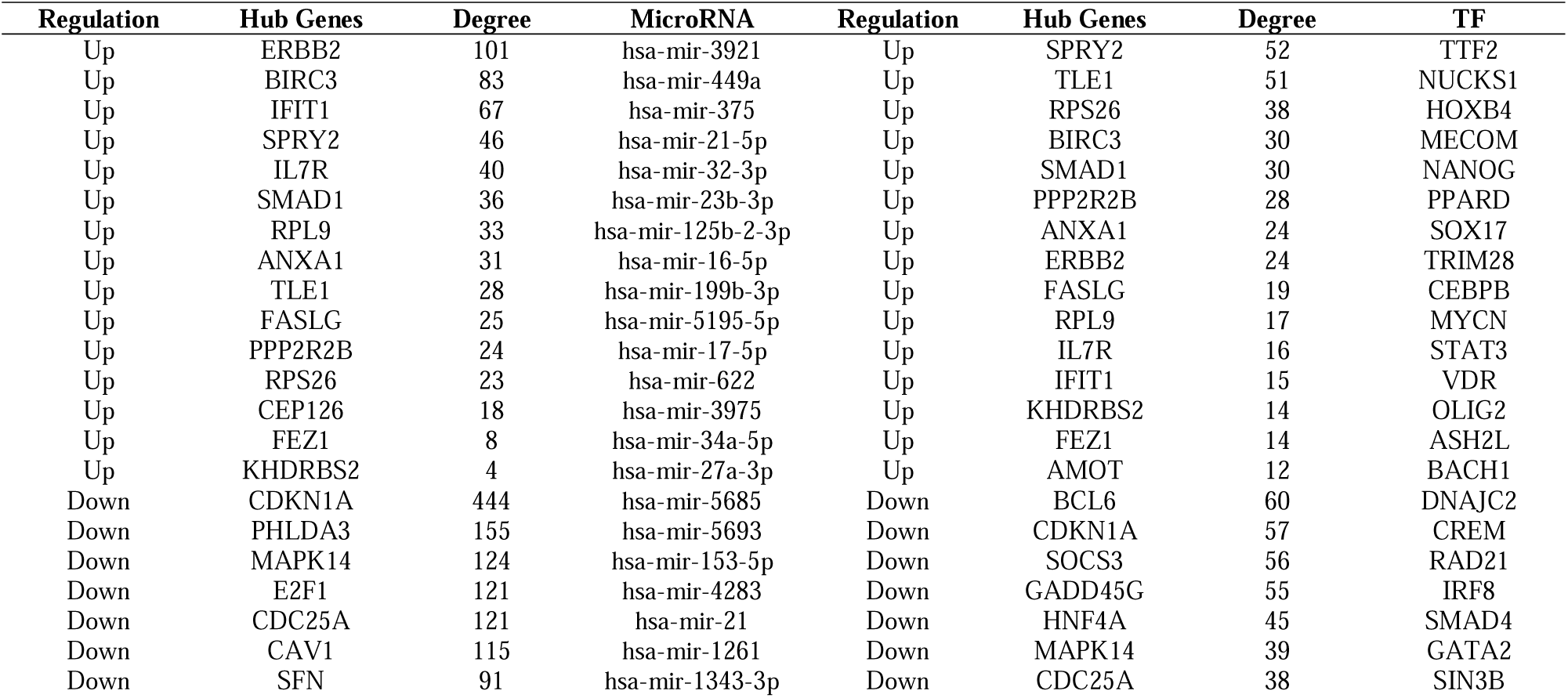
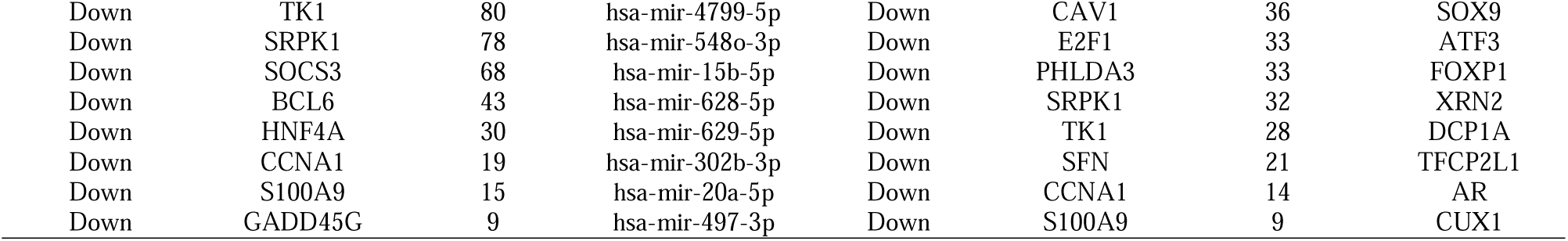
MiRNA hub gene and TF – hub gene topology table

### Construction of the TF-hub gene regulatory network

The TF-hub gene regulatory networks of the hub genes in IDB were constructed through NetworkAnalyst database, and 5745 edges involving 456 nodes (TFs: 192; Hub Genes: 264) were formed in the network (Fig. 6). SPRY2 was regulated by 52 different TFs (ex: TTF2); TLE1 was regulated by 51 different TFs (ex: NUCKS1); RPS26 was regulated by 38 different TFs (ex: HOXB4); BIRC3 was regulated by 30 different TFs (ex: MECOM); SMAD1 was regulated by 30 different TFs (ex: NANOG); BCL6 was regulated by 60 different TFs (ex: DNAJC2); CDKN1A was regulated by 57 different TFs (ex: CREM); SOCS3 was regulated by 56 different TFs (ex: RAD21); GADD45G was regulated by 55 different TFs (ex: IRF8); HNF4A was regulated by 45 different TFs (ex: SMAD4) (Table 5).

**Fig. 6.**
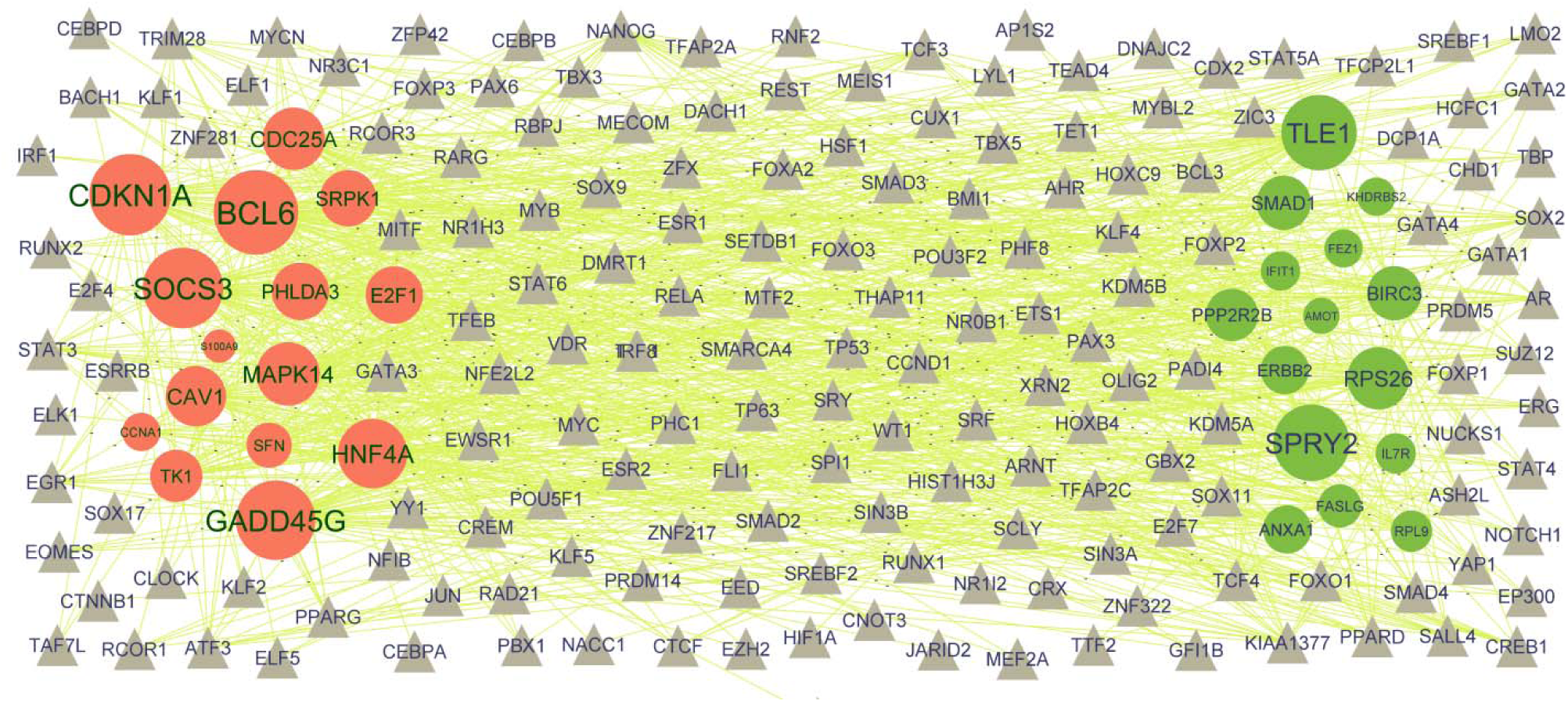
Target gene TF regulatory network between target genes. The ash color triangle nodes represent the ke TFs; up regulated genes are marked in green; down regulated genes are marked in red.

### Receiver operating characteristic curve (ROC) analysis

Furthermore, we analyzed the performance of expressed hub genes in diagnosing IBD by means of ROC curve analysis in pROC package in R statistical software. The AUC was calculated to indicate diagnostic efficiency and predictive accuracy. These expressed hub genes have superior diagnostic value in the IBD samples compared with normal control samples. Specifically, IL7R (AUC: 0.922), ERBB2 (AUC: 0.938), SMAD1 (AUC: 0.902), RPS26 (AUC: 0.915), TLE1 (AUC: 0.927), HNF4A (AUC: 0.917), CDKN1A (AUC: 0.930), SRPK1 (AUC: 0.907), H3C12 (AUC: 0.910) and SFN (AUC: 0.925) showed the highest diagnostic performance in the IBD samples (Fig.7). Based on the above data, these expressed hub genes might serve as potential biomarkers for IBD diagnosis.

**Fig. 7.**
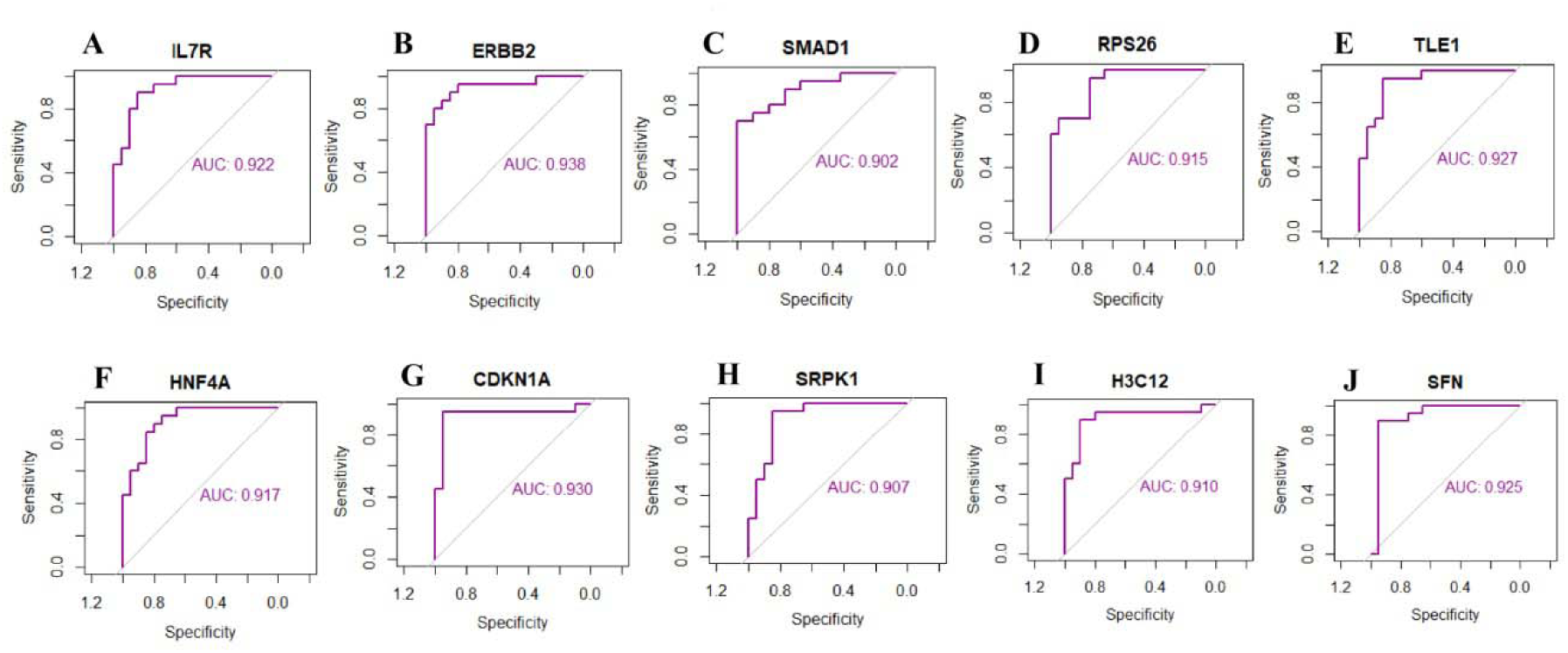
ROC curve analyses of hub genes. A) IL7R B) ERBB2 C) SMAD1 D) RPS26 E) TLE1 F) HNF4A G) CDKN1A H) SRPK1 I) H3C12 J) SFN

## Discussion

IBD is the leading cause of chronic digestive disorders globally [47]. Due to the extremely complex inflammatory disorders that occur in patients with IBD, and it is often more difficult to treat than other digestive disorders. Although numerous studies have investigated the molecular pathogenesis of IBD, it has not been clarified completely [48]. Therefore, it is necessary to identify key biomarkers for early diagnosis, prognosis, and targeted therapy of IBD. This investigation used NGS data to conduct bioinformatics analysis for identifying target genes and signaling pathways involved in the occurrence and advancement of IBD. The results of this investigation suggest that novel biomarkers might play a key role in the pathogenesis of IBD.

After the rapid advancements over the past decade, NGS has penetrated into all fields of life science and has managed to not only effectively cause advancements in research, but has also gradually been enforced in clinical practice. NGS data analysis can identify various differentially expressed genes in a short duration. NGS with bioinformatics analysis, it can help to explore disease-related biomarkers and mechanisms to predict molecular targets to develop precision therapies [49]. Therefore, bioinformatics methods in conjunction with NGS data analysis were used in our investigations to screen differential genes.

In the present investigation, we identified total 957 DEGs, including 478 up regulated genes and 479 down regulated genes. 807 IBD patients and 209 normal participants were enrolled in the current investigation. ADAM23 [50], PGF (placental growth factor) [51], CD177 [52], ANOS1 [53], RAP1GAP [54] and PRDM14 [55] are associated with the prognosis of gastrointestinal cancers. CD177 [56] was reported to be associated with the prognosis of IBD. SLC6A19 [57] has been identified as a key gene in autoimmune diseases. These findings suggested that they are positively linked with inflammatory process. These DEGs might be vital for the advancement of IBD.

Establishing GO and pathway enrichment analyses is friendly for researchers to investigate the underlying molecular mechanism of IDB for the reason that the DEGs would be ordered in the GO terms and pathways. Signaling pathways include GPCR ligand binding [58], signaling by GPCR [59], signal transduction [60], GPCR downstream signaling [61], immune system [62], hemostasis [63], neutrophil degranulation [64], infectious disease [65] and neuronal system [66] were responsible for progression of IBD. GPR15 [67], ZNF365 [68], IL2 [69], IGFBP3 [70], FASLG (Fas ligand) [71], BTNL2 [72], S100B [73], FAP (fibroblast activation protein alpha) [74], BANK1 [75], IFNG (interferon gamma) [76], ABCB1 [77], GPR183 [78], CCR4 [79], TNFRSF13C [80], FCRL3 [81], IL7R [82], ANXA1 [83], CCR6 [84], CD69 [85], ADAMDEC1 [86], HNF4A [87], SDC1 [88], MMP9 [89], HP (haptoglobin) [90], S100A12 [91], ACE2 [92], TRPV4 [93], OSM (oncostatin M) [94], ADM (adrenomedullin) [95], TLR5 [96], IGFBP2 [97], CD209 [98], SOCS3 [99], IL27 [100], S100A8 [101], AIM2 [102], GUCY2C [103], IL4R [104], S100A9 [105], CEACAM1 [106], SLC11A1 [107] and CA1 [108] are potential candidate genes for the treatment of IBD. The abnormal expression of GPR15 [109], MRGPRX2 [110], IL2 [111], IGFBP3 [112], FASLG (Fas ligand) [113], ERBB2 [114], BTNL2 [115], S100B [116], IFNG (interferon gamma) [117], ABCB1 [118], CD200 [119], FGF9 [120], CD83 [121], CD19 [122], IL5 [123], CXCR5 [124], ANXA1 [125], CCR6 [126], CCR8 [127], GZMA (granzyme A) [128], GZMB (granzyme B) [129], CD69 [130], BTLA (B and T lymphocyte associated) [131], CXCL5 [132], HNF4A [133], SDC1 [134], MMP9 [135], HP (haptoglobin) [136], GPR84 [137], S100A12 [138], CBS (cystathionine beta-synthase) [139], KCNMA1 [140], CYP1A1 [141], ACE2 [142], OSM (oncostatin M) [143], ROBO1 [144], LPO (lactoperoxidase) [145], CLDN4 [146], ADM (adrenomedullin) [147], CA3 [148], TLR5 [149], SOCS3 [150], IL27 [151], CAV1 [152], S100A8 [153], PADI4 [154], GUCY2C [155], S100A9 [156], CEACAM1 [157], SLC11A1 [158], KCNJ15 [159] and CA1 [108] might be related to the progression of ulcerative colitis. Abnormal expression of GPR15 [160], FAT4 [161], BMP3 [162], MAL2 [163], ZNF365 [164], ROR1 [165], ADAM29 [166], TAF7L [167], PAK3 [168], IL2 [169], IGFBP3 [170], FASLG (Fas ligand) [171], ERBB2 [172], WNT10A [173], WNT16 [174], ESM1 [175], PCDH17 [176], EFNA5 [177], ADAMTS1 [178], S100B [179], SORBS2 [180], CHODL (chondrolectin) [181], SPON2 [182], MACROD2 [183], MS4A1 [184], RBFOX2 [185], PLA2G4C [186], HOPX (HOP homeobox) [187], GREM2 [188], FAP (fibroblast activation protein alpha) [189], PCDHGA9 [190], AKR1C3 [191], BIRC3 [192], PDGFD (platelet derived growth factor D) [193], COL4A3 [194], IFNG (interferon gamma) [195], ABCB1 [196], AKT3 [197], CCR4 [198], ADAM28 [199], LGR6 [200], CD200 [201], TPBG (trophoblast glycoprotein) [202], ENAH (ENAH actin regulator) [203], MYO6 [204], FGF9 [205], TGFB2 [206], TRPC1 [207], EOMES (eomesodermin) [208], CD83 [209], TPD52 [210], PAX5 [211], ST8SIA6 [212], NEO1 [213], BMPR1A [214], TGFBR3 [215], AMOT (angiomotin) [216], EPHA4 [217], CAMK2N1 [218], CXCR5 [219], ABCA5 [220], DAB1 [221], SPRY2 [222], ANXA1 [223], MTSS1 [224], CCR6 [225], PTPRB (protein tyrosine phosphatase receptor type B) [226], SLC4A4 [227], PTCH1 [228], IL9R [229], PCDH9 [230], GAS1 [231], SELE (selectin E) [232], GZMA (granzyme A) [233], GZMB (granzyme B) [234], EMP1 [235], ABCB4 [236], LGALS4 [237], CD69 [238], BTLA (B and T lymphocyte associated) [239], ARHGEF28 [240], RGMB (repulsive guidance molecule BMP co-receptor b) [241], GPR63 [242], CXCL5 [243], HNF4A [244], BATF2 [245], SYT7 [246], GCKR (glucokinase regulator) [247], IGF2 [248], SDC1 [249], IGHG1 [250], MMP9 [251], PRKCG (protein kinase C gamma) [252], HP (haptoglobin) [253], GDF15 [254], GPR84 [255], S100A12 [256], ANXA3 [257], CBS (cystathionine beta-synthase) [258], BMX (BMX non-receptor tyrosine kinase) [259], NECTIN2 [260], KCNMA1 [261], NTRK3 [262], CYP1A1 [263], PLA2R1 [264], ACE2 [265], GRB10 [266], UNC5B [267], TWIST2 [268], AQP1 [269], UNC5A [270], FGF13 [271], TRPV4 [272], OSM (oncostatin M) [273], ROBO1 [274], NMU (neuromedin U) [275], BIRC5 [276], GKN2 [277], SFN (stratifin) [278], CLDN4 [279], ADM (adrenomedullin) [280], BHLHA15 [281], HPN (hepsin) [282], ITGB4 [283], GSTA1 [284], TLR5 [285], ITGA7 [286], KIF26A [287], S100P [288], IGFBP2 [289], ARG1 [290], MET (MET proto oncogene, receptor tyrosine kinase) [291], CYP2W1 [292], SOCS3 [293], CDC25A [294], CXCL2 [295], IL27 [296], BCL6 [297], PXDNL (peroxidasin like) [298], TRPM4 [299], TREM2 [300], CAV1 [301], ST3GAL4 [302], ACSL1 [303], ITLN1 [304], PROK2 [305], MAPK14 [306], CSNK1A1L [307], PADI2 [308], SFRP2 [309], MARCO (macrophage receptor with collagenous structure) [310], S100A8 [311], PDCD1LG2 [312], AIM2 [313], IQGAP3 [314], PFKFB2 [315], PADI4 [316], GJB6 [317], GUCY2C [318], GSDMC (gasdermin C) [319], IL4R [320], S100A9 [321], SEMA4A [322], PLEK2 [323], HSD11B2 [324], E2F1 [325], CDKN1A [326], SRPK1 [327], CEACAM1 [328], SLC11A1 [329], MCEMP1 [330], NRN1 [331], LRIG3 [332], INSC (INSC spindle orientation adaptor protein) [333], CD248 [334], MSLN (mesothelin) [335], TGM3 [336], FABP2 [337], SLC2A3 [338], IFITM10 [339], MUC6 [340] and CA1 [341] contributes to the gastrointestinal cancers. ZNF365 [342], IL2 [343], IGFBP3 [344], FASLG (Fas ligand) [345], BTNL2 [346], FAP (fibroblast activation protein alpha) [347], IFNG (interferon gamma) [117], ABCB1 [348], CCR4 [349], CD200 [119], FCRL3 [350], CD83 [121], CD19 [351], IL5 [352], ANXA1 [353], GZMB (granzyme B) [129], HNF4A [354], SDC1 [355], MMP9 [356], HP (haptoglobin) [357], GDF15 [358], S100A12 [359], ACE2 [360], ADM (adrenomedullin) [361], TLR5 [362], SOCS3 [363], IL27 [364], CAV1 [365], IL4R [366], S100A9 [367], E2F1 [368] and SLC11A1 [158] are important in the development of Crohn’s disease. IL2 [369] and ADAMTS1 [370] have been known to be involved in fistulas progression. Studies have reported that IL2 [371], IGFBP3 [372], IFNG (interferon gamma) [373], GDF15 [374], ACE2 [375] and IGFBP2 [376] could be related to malnutrition. IL2 [377], IGFBP3 [378], FASLG (Fas ligand) [379], BTNL2 [380], ADAMTS1 [381], S100B [382], STAP1 [383], CD160 [384], BANK1 [385], COL4A3 [386], IFNG (interferon gamma) [387], COCH (cochlin) [388], AKT3 [389], CCR4 [390], RAG1 [391], SEMA3A [392], CD200 [393], MYBL1 [394], FCRL3 [395], CD83 [396], GPR174 [397], CD19 [398], IL5 [399], EPHA4 [400], XCL1 [401], IL7R [402], CD180 [403], PRF1 [404], CXCR5 [405], ITM2A [406], ANXA1 [407], CCR6 [408], SLC4A4 [409], GZMB (granzyme B) [410], CD69 [411], BTLA (B and T lymphocyte associated) [412], IGF2 [413], SDC1 [414], MMP9 [415], GDF15 [416], C1QA [417], HRH3 [418], ACE2 [419], AQP1 [420], ROBO1 [421], SCARF1 [422], GFAP (glial fibrillary acidic protein) [423], CA3 [424], NFASC (neurofascin) [425], ADGRG3 [426], S100P [427], IGFBP2 [428], APOA2 [329], SOCS3 [430], IL27 [431], BCL6 [432], TRPM4 [433], TREM2 [434], CAV1 [435], MSR1 [436], MARCO (macrophage receptor with collagenous structure) [437], S100A8 [438], PDCD1LG2 [439], AIM2 [440], PADI4 [441], IL4R [442], S100A9 [443], SEMA4A [444], E2F1 [445], CDKN1A [446], CEACAM1 [447], SLC11A1 [448], PSTPIP2 [449] and HK3 [450] are a pathogenic genes for autoimmune diseases. Recent reports have revealed that IL2 [451] acted as biomarker in coagulation and fibrinolysis. Research has shown that IL2 [452], IGFBP3 [70], ENPP1 [453], FASLG (Fas ligand) [454], FAP (fibroblast activation protein alpha) [455], IFNG (interferon gamma) [456], AKT3 [457], BMPR1A [458], SPRY1 [459], ANXA1 [460], PTCH1 [461], IGF2 [462], MMP9 [463], HP (haptoglobin) [464], GDF15 [465], KCNMA1 [466], ACE2 [467], TRPV4 [468], ADM (adrenomedullin) [469], CD274 [470], IGFBP2 [471], SOCS3 [472], TREM2 [473], CAV1 [474], PROK2 [475], MAPK14 [476] and CDKN1A [477] plays an important role in the pathogenesis of osteoporosis. Studies have revealed that IL2 [478], ENPP1 [479], FASLG (Fas ligand) [480], FAP (fibroblast activation protein alpha) [481], BIRC3 [482], IFNG (interferon gamma) [483], NCR1 [484], RAG1 [485], SEMA3A [486], TIGIT (T cell immunoreceptor with Ig and ITIM domains) [487], CD19 [488], TGFBR3 [489], IL5 [490], MMP9 [491], HP (haptoglobin) [492], GDF15 [493], ACE2 [494], DYSF (dysferlin) [495], IL27 [496], CAV1 [497], ALAS2 [498], S100A8 [499] and S100A9 [499] plays a key role in anemia. Studies had shown that IGFBP3 [344] were associated with intestinal strictures. Therefore, these GO terms and signaling pathways are most likely to be important in the development of IDB and IDB complications. Additional investigations are required to identify all the DEGs in IDB.

To screen potential diagnostic and prognostic biomarkers for IBD, we performed PPI network construction and module analysis by the use of the above DEGs, and only hub genes were identified. IL7R [82] and HNF4A [87] plays a crucial role in the pathogenesis of IBD and it is also considered a promising target for pharmacological-based therapies. IL7R [402] and CDKN1A [446] are an essential mediator of inflammation, which is closely related to the progression of autoimmune diseases. ERBB2 [114] and HNF4A [133] are reported to play a central role in ulcerative colitis. It was reported that ERBB2 [172], HNF4A [244], CDKN1A [326], SRPK1 [327], SFN (stratifin) [278] and RPL9 [500] participate in pathogenic processes of gastrointestinal cancers. A study indicates that RPS26 [501] is positively correlated with anemia. Studies had shown that TLE1 [502] and HNF4A [354] were associated with Crohn’s disease. CDKN1A [477] plays an indispensable role in osteoporosis. Our findings suggested SMAD1, H3C12 and GADD45G as potential novel diagnostic biomarkers for IBD. Therefore, our study may contribute to the understanding of the molecular mechanisms affecting inflamation after IDB.

MiRNA-hub gene regulatory network and TF-hub gene regulatory network were successfully constructed via the miRNet and NetworkAnalyst online database and cytoscape software. Vital regulated hub genes, miRNA and TFs were screened from the MiRNA-hub gene regulatory network and TF-hub gene regulatory network. ERBB2 [114], CAV1 [152], SOCS3 [150], HNF4A [133], hsa-mir-449a [503], hsa-mir-21 [504], NUCKS1 [505], NANOG (Nanog homeobox) [506], IRF8 [507] and SMAD4 [508] were a diagnostic markers of ulcerative colitis and could be used as therapeutic targets. ERBB2 [172], BIRC3 [192], IFIT1 [509], SPRY2 [222], MAPK14 [306], E2F1 [325], CDC25A [294], CAV1 [301], BCL6 [297], CDKN1A [326], SOCS3 [293], HNF4A [244], hsa-mir-449a [503], hsa-mir-375 [510], hsa-mir-21-5p [511], hsa-mir-21 [512], NUCKS1 [513], HOXB4 [514], MECOM (MDS1 and EVI1 complex locus) [515], NANOG (Nanog homeobox) [506], DNAJC2 [516], RAD21 [517], IRF8 [507] and SMAD4 [518] were found to be substantially related to gastrointestinal cancers. BIRC3 [482], RPS26 [501], HOXB4 [519] and SMAD4 [520] might be associated with the prognosis of anemia. IL7R [82], SOCS3 [99], HNF4A [87], hsa-mir-375 [521], hsa-mir-21 [522], MECOM (MDS1 and EVI1 complex locus) [515] and CREM (cAMP responsive element modulator) [523] are a diagnostic biomarkers and immunotherapeutic target for IBD. Studies have found that IL7R [402], E2F1 [445], CAV1 [435], BCL6 [432], CDKN1A [446], SOCS3 [430], IRF8 [524] and SMAD4 [525] might be a prognostic biomarker and potential therapeutic target for patients with autoimmune diseases. MAPK14 [476], CAV1 [474], CDKN1A [477], SOCS3 [472], hsa-mir-21-5p [526] and SMAD4 [527] can be used as a diagnostic marker for osteoporosis. E2F1 [368], CAV1 [365], TLE1 [502], SOCS3 [363], HNF4A [354], hsa-mir-21 [528] and SMAD4 [529] contributes to the progression of Crohn’s disease. PHLDA3, SMAD1, GADD45G, hsa-mir-3921, hsa-mir-32-3p, hsa-mir-5685, hsa-mir-5693, hsa-mir-153-5p, hsa-mir-4283 and TTF2 might be novel therapeutic targets for IDB. Therefore, all the hub genes, miRNA and TFs may possess key roles in IDB and could interact with each other. They might be used as potential effective candidates for early diagnosis or prognosis.

In conclusion, the present investigation provided a pipeline for the analysis of NGS dataset. In addition, numerous known and potential novel markers were proposed in the present investigation, which might contribute to diagnosis and treatment targets for IDB; however, further investigation is required for confirmation of the functions exhibited by these biomarkers.

## Acknowledgement

I Carmen argmann, MSSM, new york city, USA, very much, the author who deposited their NGS dataset GSE186507, into the public GEO database.

## Ethical approval

This article does not contain any studies with human participants or animals performed by any of the authors.

## Informed consent

No informed consent because this study does not contain human or animals participants.

## Availability of data and materials

The datasets supporting the conclusions of this article are available in the GEO (Gene Expression Omnibus) (https://www.ncbi.nlm.nih.gov/geo/) repository. [(GSE186507) https://www.ncbi.nlm.nih.gov/geo/query/acc.cgi]

## Consent for publication

Not applicable.

## Author Contributions

B. V. - Writing original draft, and review and editing

C. V. - Software and investigation

